# Single Cell RNA-seq reveals ectopic and aberrant lung resident cell populations in Idiopathic Pulmonary Fibrosis

**DOI:** 10.1101/759902

**Authors:** Taylor S. Adams, Jonas C. Schupp, Sergio Poli, Ehab A. Ayaub, Nir Neumark, Farida Ahangari, Sarah G. Chu, Benjamin A. Raby, Giuseppe DeIuliis, Michael Januszyk, Qiaonan Duan, Heather A. Arnett, Asim Siddiqui, George R. Washko, Robert Homer, Xiting Yan, Ivan O. Rosas, Naftali Kaminski

## Abstract

We provide a single cell atlas of Idiopathic Pulmonary Fibrosis (IPF), a fatal interstitial lung disease, focusing on resident lung cell populations. By profiling 312,928 cells from 32 IPF, 29 healthy control and 18 chronic obstructive pulmonary disease (COPD) lungs, we demonstrate that IPF is characterized by changes in discrete subpopulations of cells in the three major parenchymal compartments: the epithelium, endothelium and stroma. Among epithelial cells, we identify a novel population of IPF enriched aberrant basaloid cells that co-express basal epithelial markers, mesenchymal markers, senescence markers, developmental transcription factors and are located at the edge of myofibroblast foci in the IPF lung. Among vascular endothelial cells in the in IPF lung parenchyma we identify an expanded cell population transcriptomically identical to vascular endothelial cells normally restricted to the bronchial circulation. We confirm the presence of both populations by immunohistochemistry and independent datasets. Among stromal cells we identify fibroblasts and myofibroblasts in both control and IPF lungs and leverage manifold-based algorithms diffusion maps and diffusion pseudotime to infer the origins of the activated IPF myofibroblast. Our work provides a comprehensive catalogue of the aberrant cellular transcriptional programs in IPF, demonstrates a new framework for analyzing complex disease with scRNAseq, and provides the largest lung disease single-cell atlas to date.

## Main Text

Idiopathic Pulmonary Fibrosis (IPF) is a progressive lung disease characterized by irreversible scarring of the distal lung, leading to respiratory failure and death (*1, 2*). Despite significant progress in our understanding of pulmonary fibrosis in laboratory animals, we have a limited perspective of the cellular and molecular processes that determine the IPF lung phenotype. In fact, IPF is still best described by its histopathological pattern of usual interstitial pneumonia (UIP), that includes presence of fibroblast foci; hyperplastic alveolar epithelial cells that localize adjacent to fibroblastic foci; a distortion of airway architecture combined with an accumulation of microscopic airway-epithelial lined cysts known as “honeycombs” in the distal parenchyma and the lack of evidence for other conditions(*1-3*).

Evidence for molecular aberrations in the IPF lung have mostly been obtained by following hypotheses derived from animal models of disease, from discovery of genetic associations in humans, or from genes differentially expressed in transcriptomic studies of bulk IPF tissue with limited cellular resolution (*4, 5*). Recent studies have demonstrated the value of single cell RNA sequencing (scRNAseq) by identifying profibrotic macrophages in lungs of human and mice with pulmonary fibrosis (*6, 7*). Here we harness the cell-level resolution afforded by scRNAseq to provide an atlas of the extent of complexity and diversity of aberrant cellular populations in the three major parenchymal compartments of the IPF lung: the epithelium, endothelium and stroma.

## Results

We profiled 312,928 cells from distal lung parenchyma samples obtained from 32 IPF, 18 COPD and 29 control donor lungs (Table S1 and Fig. 1A), identifying 38 discrete cell types (Fig. 1B) based on distinct markers (Figure 1C, Data S1-S4). Manually curated cell classifications are consistent with automated annotations drawn from several independent databases (Fig. S4). The detailed cellular repertoires of epithelial, endothelial and mesenchymal cells are provided below. Our data will be available on GEO (GSE136831) and the results can be explored through the IPF Cell Atlas Data Mining Site (*8*).

**Fig. 1.**
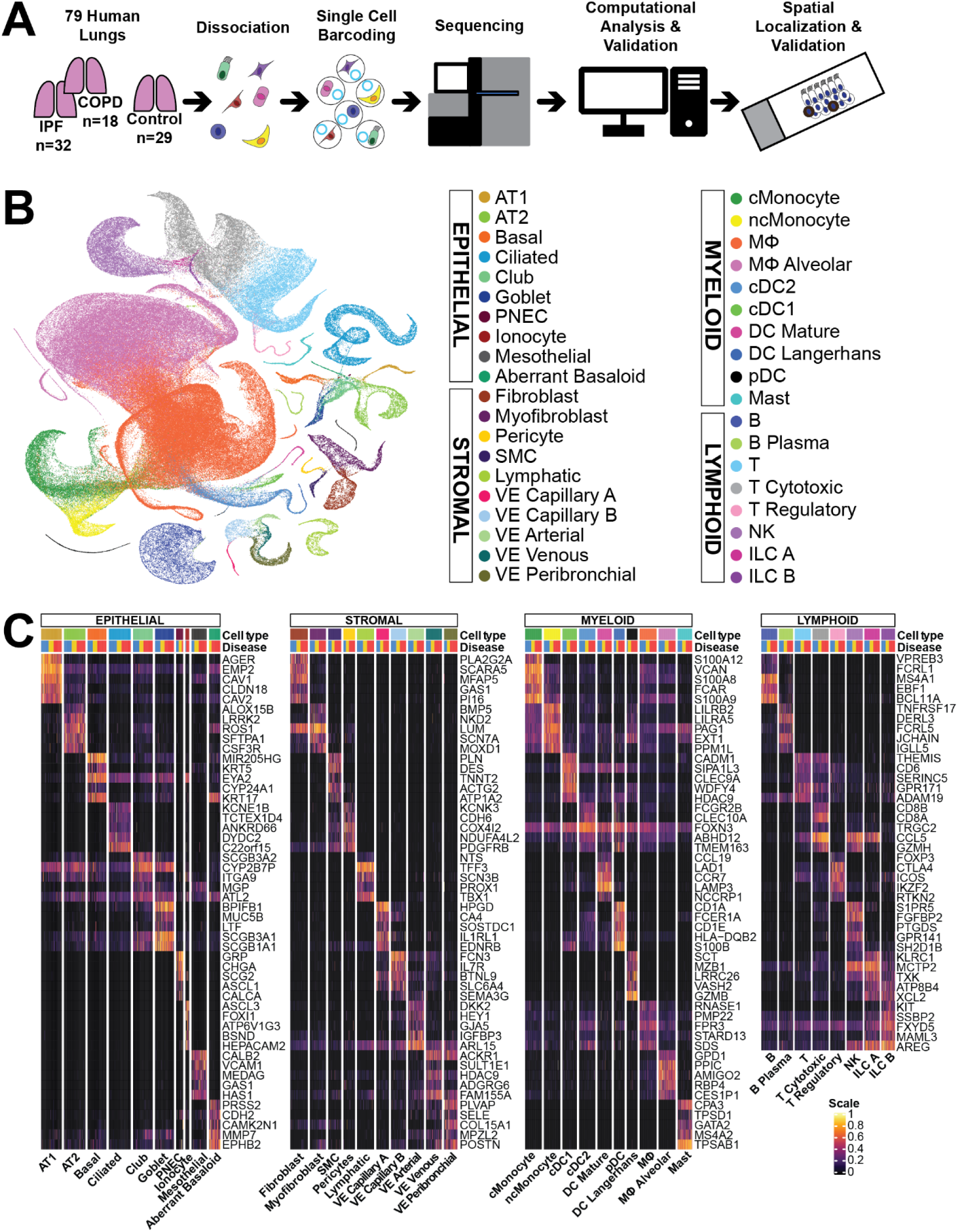
Profiling Human Lung Heterogeneity with scRNAseq. **(A)** Overview of experimental design. (i) Disease lung explants and unused donor lungs collected (ii) Lungs dissociated to single cell suspension (iii) Droplet-based scRNAseq library prep (iv) sequencing (v) Exploratory analysis (vi) Spatial localization with IHC. **(B)** UMAP representation of 312,928 cells from 32 IPF, 18 COPD and 29 control donor lungs; each dot represents a single cell, and cells are labelled as one of 38 discrete cell varieties. **(C**) Heatmap of marker genes for all 38 identified cell types, categorized into 4 broad cell categories. Each cell type is represented by the top 5 genes ranked by false-detection-rate adjusted p-value of a Wilcoxon rank-sum test between the average expression per subject value for each cell type against the other average subject expression of the other cell types in their respective grouping. Each column represents the average expression value for one subject, hierarchically grouped by disease status and cell type. Gene expression values are unity normalized from 0 to 1.

### The epithelial cell repertoire of the fibrotic lung is dramatically changed and contains disease associated aberrant basaloid cells

In non-diseased tissue we identified all known lung epithelial cells populations including, alveolar type 1 and type 2 cells, ciliated cells, basal cells, goblet cells, club cells, pulmonary neuroendocrine cells and ionocytes (Fig. 1B-C). The epithelial cell repertoire of IPF lungs is characterized by an increased proportion of airway epithelial cells and substantial decline in alveolar epithelial cells (Fig. 2A-B, Data S6). Impressively, nearly every epithelial cell type we identified exhibited profound changes in gene expression in the IPF lung compared to control or COPD (Data S7, S9; IPF cell Atlas Data Mining Site (*8*)). Among epithelial cells we identified a population of cells that was transcriptionally distinct from of any epithelial cell type previously described in the lung that we termed aberrant basaloid cells (Fig. 2C). In addition to epithelial markers, these cells express the basal cell markers TP63, KRT17, LAMB3 and LAMC2, but do not express other established basal markers such as KRT5 and KRT15. These cells express markers of epithelial mesenchymal transition (EMT) such as VIM, CDH2, FN1, COL1A1, TNC, and HMGA2 (*9, 10, 11, 12*), senescence related genes including CDKN1A, CDKN2A, CCND1, CCND2, MDM2, and GDF15 (*13-15*) (Figure 2C, right) and the highest levels of established IPF-related molecules, such as MMP7 (*16*), αVβ6 subunits (*17*) and EPHB2 (*18*). The cells exhibit high levels of expression and evidence of downstream activation of SOX9 (Figure 2C upper panel), a transcription factor critical to distal airway development, repair, also indicated in oncogenesis (*19, 20*). No aberrant basaloid cells could be detected in control lungs (Fig. 2A-B). Of the 483 aberrant basaloid cells identified, 448 were from IPF lungs compared to just 33 cells from COPD lungs. We confirmed the presence and localized these cells in the IPF lung by immunohistochemistry (IHC) using p63, KRT17, HMGA2, COX2 and p21 as markers (Fig. 2D). In IPF lungs, these cells consistently localize to the epithelial layer covering myofibroblast foci. No aberrant basaloid could be identified in control lungs. To independently validate our results, we reanalyzed the IPF single cell data published by Reyfman *et al*.*(6)*. We identified 80 aberrant basaloid cells in this data and observed greater correlation between them from each dataset (Spearman’s rho: 0.81), than with epithelial cells within each dataset (Figure 2E) confirming our results.

**Fig. 2.**
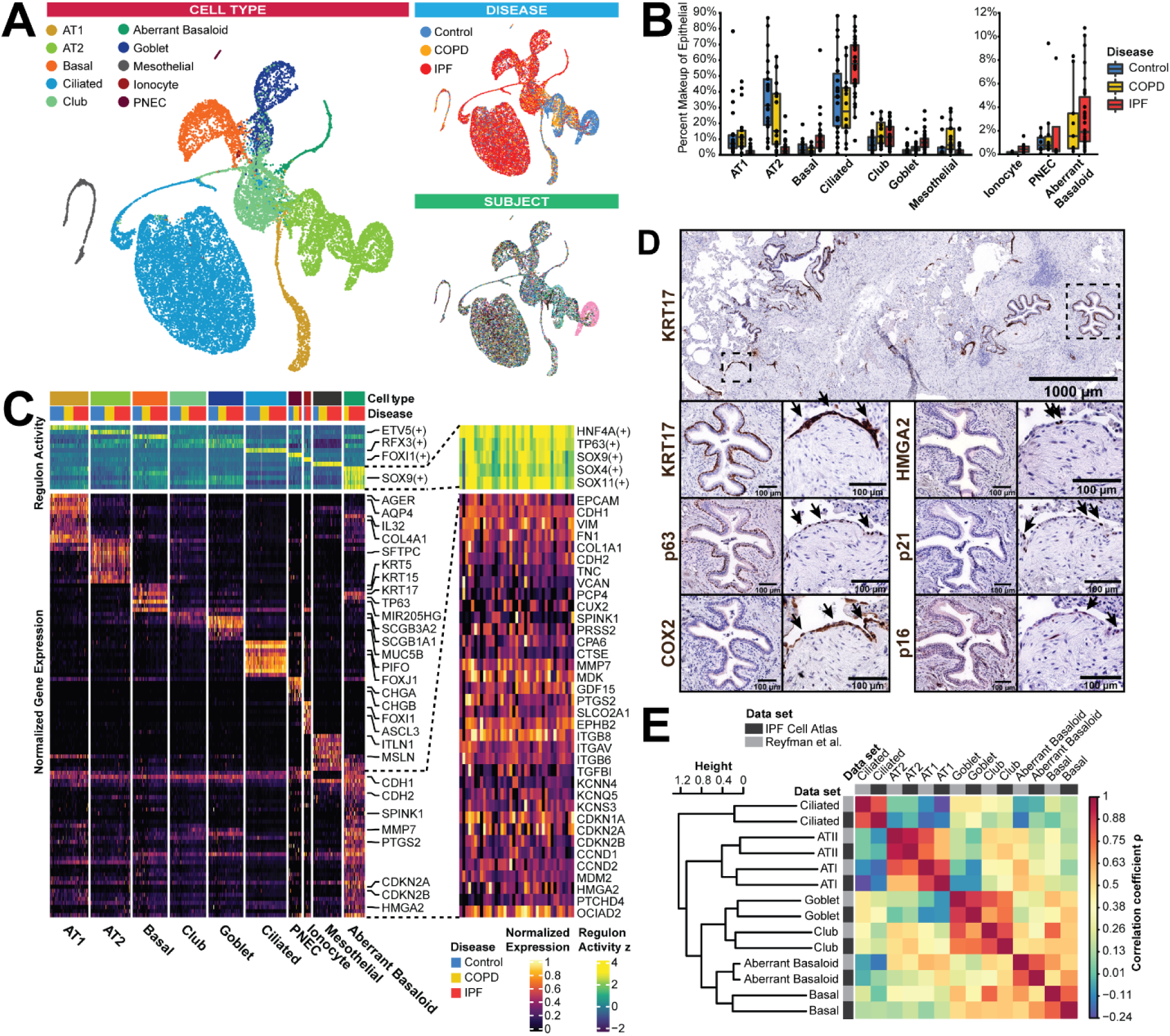
Identification of Aberrant Basaloid Cells in IPF and COPD Lungs. **(A)** UMAPs of all epithelial cells labeled by cell type (upper left), disease status (upper right) and subject (lower right). In subject plot, each color depicts a distinct subject. **(B)** Boxplots representing the percent makeup distributions of epithelial cell types as a proportion of all sampled epithelial cells per subject, within each disease group. Each dot represents a single subject, whiskers represent 1.5 x interquartile range (IQR). **(C)** Heatmap of average gene expression and predicted transcription factor activity per subject across each of the identified epithelial cell types. Columns are hierarchically ordered by disease status and cell type, with the above annotation bar representative of the disease status. Upper green: Predicted transcription factor signatures. Right: Zoom annotation of distinguishing markers for aberrant basaloid cells. **(D)** Immunohistochemistry staining of aberrant basaloid cells in IPF lungs: epithelial cells covering fibroblast foci are p63+KRT17+ basaloid cells staining COX2, p21 and HMGA2 positive, while basal cells in bronchi do not. **(E)** Correlation matrix of epithelial cell populations we identified in independent dataset *(6)* with analogous cell types from our data. Matrix cells are colored by Spearman’s rho, cell populations are ordered with hierarchical clustering. The origin dataset for each cell population is denoted by in the annotation bars.

### The endothelial cell repertoire of the IPF lung contains ectopic peribronchial endothelial cells

While changes in vascular endothelium have been long noted in the IPF lung, little is known about their cellular and molecular characteristics (*21*). Cluster analysis of vascular endothelial (VE) cells reveal 4 populations readily characterized as capillary, arterial or venous VE cells (Fig. 3A-C). A fifth VE population is best distinguished by its expression of COL15A1. IHC assessment of COL15A1 in healthy lungs obtained from the Human Protein Atlas (*22*) revealed that COL15A1+ VE cells are restricted to vasculature adjacent to major airways (Fig. S7); we consequently refer to these cells as peribronchial VE (pVE). While pVE cells are found in subjects from all disease states, they are substantially more abundant in IPF lungs compared to controls or COPD (median percentage of VE: 54%, 8.9%, 7.1% respectively; Fig. 3C). Localization of pVE cells using the pan-endothelial marker CD31 alongside COL15A1 confirms that within control lungs, they are indeed confined to bronchial vasculature surrounding large proximal airways (Fig. 3D). In IPF lungs, COL15A1+ CD31+ VE cells are observed in the distal lung, typically at the parenchymal edge of fibroblastic foci and in areas of bronchiolization (Fig 3D). Re-analysis of a recently published scRNAseq dataset that contained normal airway and lung parenchyma samples (*23*) confirmed that genes specific to the pVE were not observed in distal lung VE (Fig. 3E). Collectively, these observations indicate that COL15A1+ VE cells, represent an ectopic pVE population in the distal lung in IPF.

**Fig. 3.**
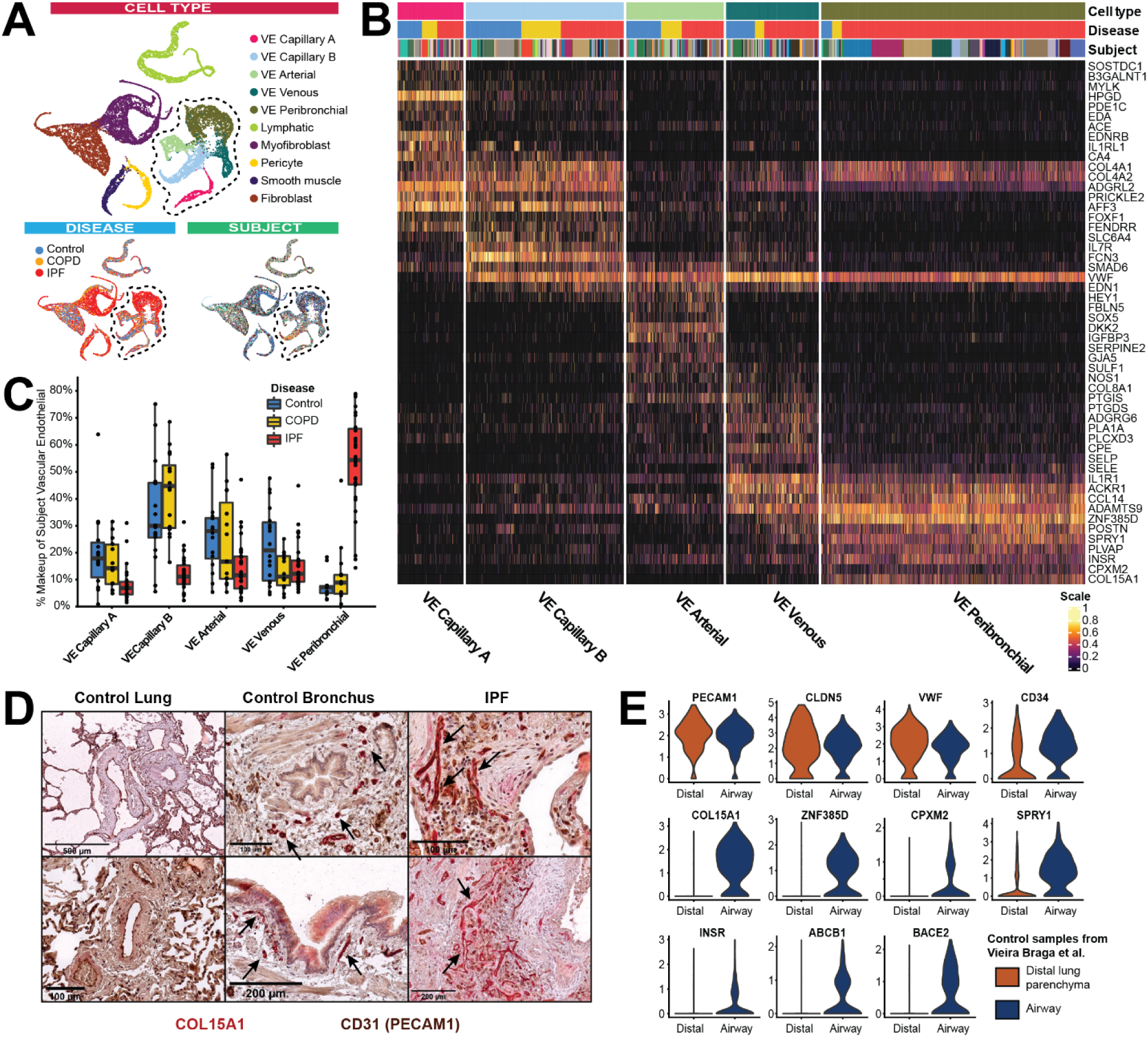
Identification of Disease-Enriched Vascular Endothelial Cell Population. **(A)** UMAPs of endothelial and mesenchymal cells labelled by cell type, disease status and subject. In subject plot, each color represented by a unique color. **(B)** Heatmap representing characteristics of 5 sub types of VE cells. Gene expression is unity normalized between 0 and 1. Each column represents an individual cell, information regarding subject, disease state, then VE type is represented in the colored annotation bars above. Each subject is represented by a unique color. **(C)** Boxplots representing the percent makeup distributions of each VE cell-type amongst all VE cells, within each disease group. Each dot represents a single subject, whiskers represent 1.5x IQR. **(D)** Immunohistochemistry staining for CD31 (PECAM1) and COL15A1 in control distal lung, control proximal lung, and affected regions of distal IPF lungs. **(E)** Violin plots of expression of pan-VE markers and peribronchial VE-specific markers across VE cells from distal and airway lung samples, from independent dataset (*23*).

### The IPF lung exhibits disease-specific archetypes among fibroblasts and myofibroblast

To characterize myofibroblasts and fibroblasts in the IPF lung, we first focused on cell populations characterized by PDGFRB expression, we then removed cells characterized as smooth muscle cells (DES, ACTG2 and PLN) or pericytes (RGS5, COX4I2). This strategy allowed us to identify two distinct stromal populations: fibroblasts, characterized by expression of CD34, FBN1, FBLN2 and VIT; myofibroblasts consistently express MYLK, NEBL, MYO10, MYO1D, RYR2 and ITGA8, as described in the murine lung (*24*) (Fig. 4A). While these features are consistent across both cell populations in IPF and control lungs (Fig. 4A-B), the IPF lung contains a myofibroblast phenotype enriched with collagens, COL8A1 and ACTA2 (Fig. 3B-D) and a fibroblast phenotype that exhibits increased expression of HAS1, HAS2, FBN1 and the SHH induced chemokine, CXCL14 (*25*) (Fig. 3B-D). Application of lineage reconstruction technique partition based graph abstraction (PAGA) (*26*) to subclustered population of both cell types (Fig. 4B) reveal that the likelihood of connective structures in phenotype space between fibroblast and myofibroblast is relatively weak (edge confidence < 0.2), when compared to connectivity within either fibroblast or myofibroblast subpopulations, irrespective of disease state enrichment (edge confidence between 0.3 and 0.6) (Fig. 4B lower). Characterization of the extent of cellular diversity along the phenotypic manifold ranging from control-enriched quiescence to disease-enriched archetype in both myofibroblast and fibroblast populations identified disease archetype changes in gene expression, with genes commonly associated with invasive fibroblasts or activated myofibroblasts at the IPF edge of the manifold (Fig. 4C-E). Taken together, these results suggest a continuous trajectory within fibroblast and myofibroblast, and not across them.

**Fig 4.**
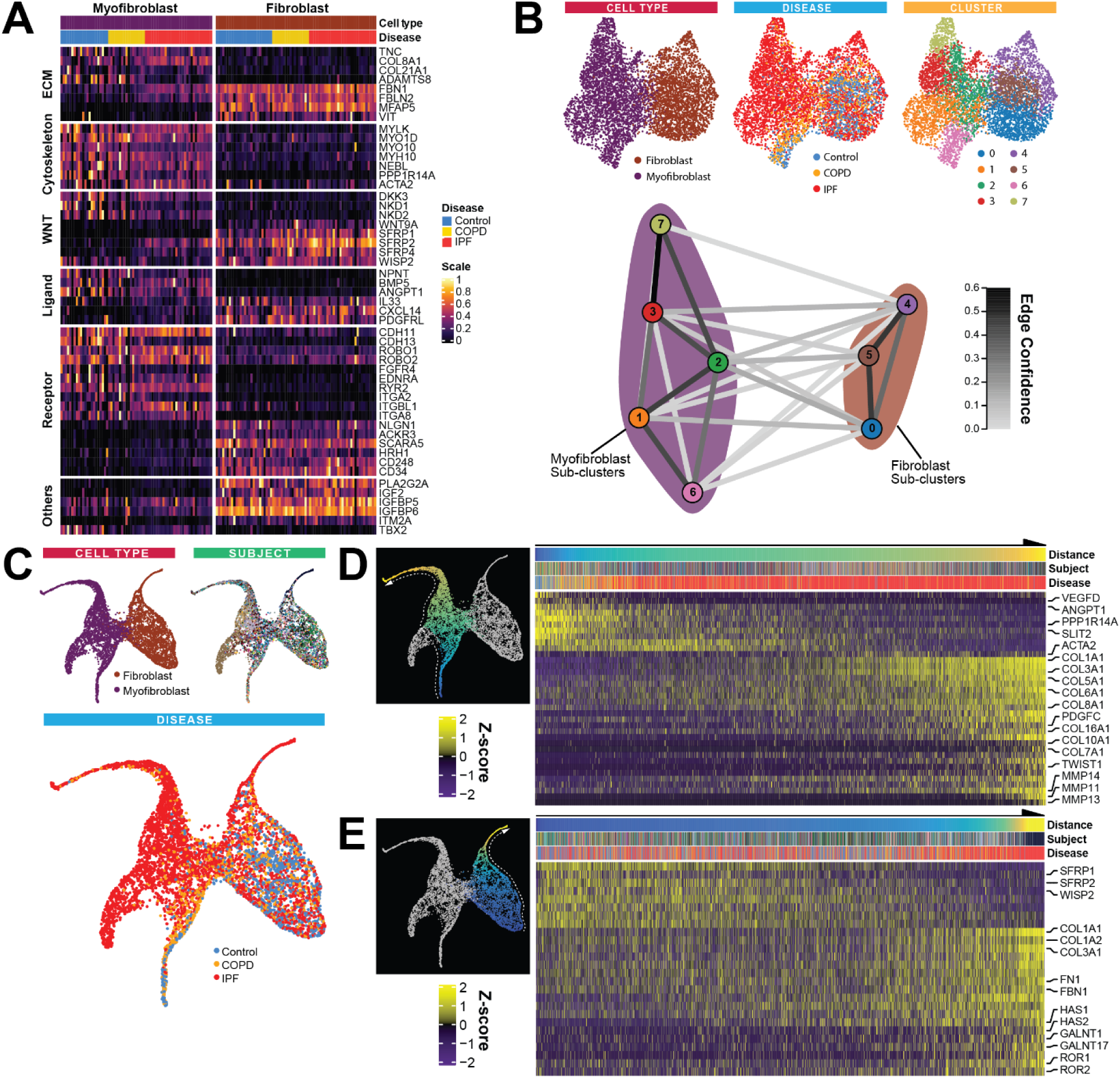
IPF Fibroblast and Myofibroblast Archetype Analysis. **(A)** Heatmap of unity-normalized gene expression of markers that delineate myofibroblast and fibroblast; each column is representative of the average expression value per cell-type for one subject. **(B)** Above: UMAPs of myofibroblast and fibroblast labelled by cell type, disease and unsupervised Louvain sub-clusters. Below: Partition graph abstraction (PAGA) analysis. Nodes represent sub-clusters; edges represent the probability of inter-node overlap based on the underlying network of cell neighborhoods. **(C)** UMAPs of myofibroblast and fibroblast cells following diffusion map implementation labelled by cell type, disease status and subject. In subject plot, each color represents a unique subject. **(D)** and **(E)** Heatmaps of myofibroblast and fibroblast respectively, ordered by diffusion pseudo time (DPT) distances along UMAP manifolds that transition from control-enriched regions towards IPF-enriched archetypes, left to right; annotation bars represent the distance, subject and disease status for each cell. Expression values are centered and scaled.

### The IPF lung gene regulatory network deviates markedly from homeostatic condition

To better understand how lung global regulatory networks were altered in IPF we implemented the gene regulatory network (GRN) inference approach BigSCale (*27*) to control and IPF cells (Fig. 5A). The topology of the IPF GRN exhibited higher density and modularity (Fig. 5B). Comparing the array of cellular contributions to communities that comprise each GRN, we found that control GRN communities show a relatively diverse array of cellular contributions both within communities and across them - sconsistent with organ function under homeostatic conditions. In IPF GRN, the major epithelial-associated communities remain largely isolated from the rest of the network, whose community with the highest density is predominantly driven by aberrant basaloid cells (Fig. 5C). Using PageRank (*28*) centrality as a proxy for a gene’s influence on the network, we identified the top 300 influencer genes ranked by PageRank differential compared to the control GRN (Fig. 5D). Gene set enrichment of these genes returned cellular aging, senescence, response to TGFB1, epithelial tube formation and smooth muscle cell differentiation (Figure5E). These findings underscore the heterotypic cellular contributions to the many key processes of molecular aberrancies in IPF.

**Fig. 5.**
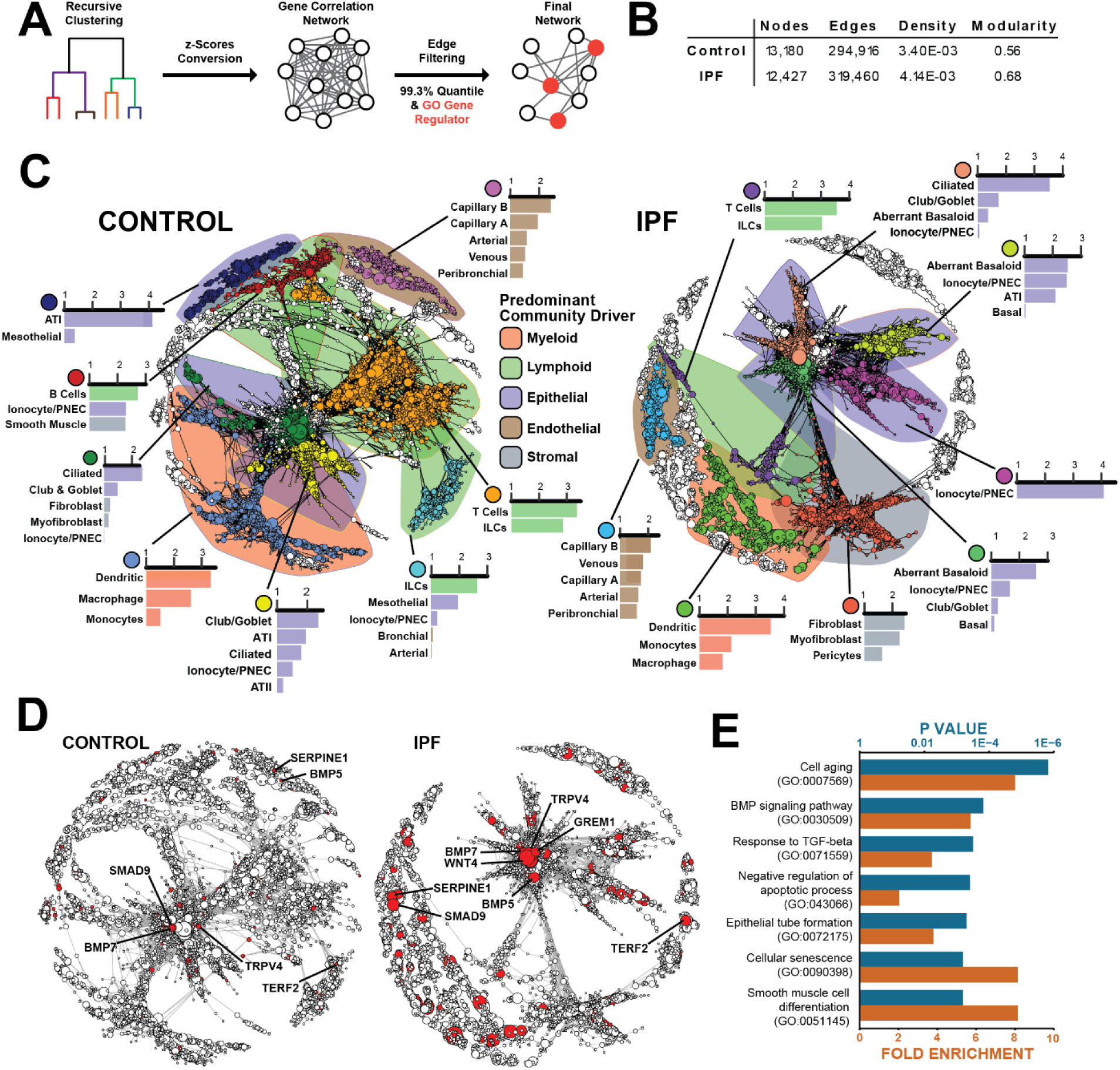
Gene Regulatory Network Analysis. **(A)** Overview of bigSCale method for computing gene regulatory network (i) cells are recursively clustered down to sub-clusters (ii) z-scores are calculated based on differential expression between sub-clusters (iii) correlations between all genes via Pearson and cosine (iv) correlation edges are thresholded using the top 99.3% quantile of correlation coefficients; only edges where at least one node has a GO annotation as a gene regulator **(B)** Summary of network structure for control and IPF GRNs. **(C)** Gene regulatory networks of control and IPF cells. Nodes represent genes, edges represent correlations of putative regulatory relationships. Nodes sizes correspond to PageRank centralities, the largest clusters are assigned colors to their nodes, with each color representing a distinct cluster. The top cell types relevant to each highlighted cluster are shown. Behind each highlighted cluster is a polygon shape covering the domain of the cluster, colored by the category of cell type that’s predominantly relevant to the community. **(D)** The same GRNs with the top 300 nodes ranked by differential PageRank centrality between IPF and control highlighted in red. Nodes sizes correspond to PageRank centralities. **(E)** Selected results from GO gene set enrichment of the top 300 differential PageRank nodes between IPF and controls, with all nodes used as a reference.

## Discussion

In this study, we provide a single cell atlas of the IPF lung, with a focus on aberrant epithelial, fibroblast and endothelial cell populations. Among epithelial cells, we discover a shift in epithelial cell population composition in the distal lung, from alveolar epithelial cells towards enrichment in cell populations that typically reside in the airway. We identify the existence of aberrant basaloid cells: a rare, disease-enriched and novel epithelial cell population that co-express basal epithelial markers, mesenchymal markers, senescence markers, developmental transcription factors and known markers of IPF, and are located at the edge of myofibroblast foci in the IPF lung. Analysis of vascular endothelial cells reveals an expanded VE population expressing markers usually characteristic of VE cells restricted to the bronchial circulation. This population localizes to areas of remodeling and aberrant angiogenesis in the distal lung parenchyma in IPF, is restricted to the airways in the normal lung, and represents a novel cellular aberration in IPF. Among stromal cells we identify two independent fibrotic archetypes associated with stromal populations present in the normal lung. Invasive fibroblasts likely related to resident lung fibroblasts and IPF myofibroblasts related to resident normal lung myofibroblasts. The extent of the impact of aberrant cell populations on the phenotype of the IPF lung is highlighted by our gene regulatory network analysis that reveals a shift from a balanced multicellular gene regulatory network in the homeostatic lung, to a fragmented network dominated by the aberrant basaloid cells, the IPF fibroblasts and myofibroblasts.

Our results represent an important contribution to our understanding of the involvement and extent of aberrations of epithelial cells in the human IPF lung. The aberrant basaloid cell, a novel, rare population of cells that never appear in control lungs and could easily be confused with airway basal cells are of particular interest. These cells express some of the most typical airway basal cell markers but not all of them. They express EMT and senescence markers as well as molecules associated with IPF, and genes that are typically expressed in other organs. Importantly, they express SOX9, a transcription factor critical to distal airway development, repair, also indicated in oncogenesis (*19, 20*). Anatomically, these cells localize to a highly enigmatic region in the IPF lung, at the active edge of the myofibroblast foci. Thus, the identification of these cells may provide a critical answer to the question of the monolayer of cuboidal epithelial cells found on the surface of IPF myofibroblastic foci. Sometimes referred to as hyperplastic alveolar epithelial cells (*29, 30*), the nature of this cells been a source for controversy, because they could not be fully characterized with previous techniques (*30-32*). Taken together with previous observations demonstrating the presence of p63+ cells undergoing partial EMT (*33, 34*), our results imply that this population is indeed distinct from resident alveolar epithelial cell populations and perhaps derived from a rare progenitor niche with the potential to serve as secondary progenitor for depleted AT1 and AT2 cells in normal human lungs, much like broncho-alveolar stem cells are known to do in the murine lung (*35, 36*). In IPF, repeated damage and potential genetic predisposition to replicative exhaustion could perhaps lead to the conflicting state of proliferation, differentiation and senescence apparent in these aberrant basaloid cells. Speculation aside, the revelation that these cells are fundamentally basaloid in nature, as well as the expansion of regular airway basal cells in the IPF lung serves to couple the two most distinct histopathological features of IPF - fibroblastic foci and honeycomb cysts - to a singular commonality: the migration of airway basal cells from their natural airway niche into the distal lung in a failed attempt to repair the lung.

The absence of discriminating markers for lung vascular endothelial cells, and the difficulty in culturing these cells has limited investigations into vascular remodeling in IPF. Our scRNAseq analysis led to the unexpected discovery of an expanded VE cell population that expresses COL15A1 in the IPF lung. These peribronchial VE cells are transcriptomically indistinguishable from systemically supplied bronchial VE cells detected in control lungs, but they localize to the vessels underneath fibroblastic foci and honeycomb cysts in IPF. While lacking support in more recent observations, it is possible that our discovery provides the cellular molecular correlate of Turner-Warwick’s 1963 observation that the bronchial vascular network is expanded throughout the IPF lung (*37*). While it is impossible at this stage to tell whether the ectopic presence of peribronchial VE cells is involved in the pathogenesis of IPF, this novel finding fits a larger pattern in the IPF lung, where cellular population normally relegated to the airways are found in affected regions in the distal parenchyma.

While it is well recognized that lung fibroblast populations demonstrate considerable plasticity (*35, 38-42*), cell-types are traditionally defined by the presence of a singular molecular feature. One relevant example being the use of ACTA2 to define the IPF myofibroblast that comprise the disease’s lesions. The cellular source of this pathological cell population remains a matter of considerable debate, as cells that satisfy this absolute definition are not present in the normal adult lung parenchyma. scRNAseq analysis provides a more comprehensive view of cellular identity, where global transcriptomic features - rather than singular cell markers - are exploited to define cells and assess the extent of population variance. Our analysis suggests that pathological, ACTA2-expressing IPF myofibroblast are not a discrete cell-type, but rather one extreme pole of a continuum connected to a quiescent ACTA2-negative stromal population represented in control lungs. We find the cells that belong to this continuum can collectively be distinguished from other lung fibroblast by a panel of genes which include several myosin and contractile-associated features, suggesting that the pathological phenotype observed in IPF is a latent feature of this population, rather than the result of trans-differentiation from a discrete cellular population. More explicitly, we did not observe any evidence suggesting that resident lung fibroblasts or activated fibroblasts were the source for IPF myofibroblasts.

Our results provide a comprehensive portrait of the fibrotic niche in IPF: where aberrant basaloid cells interface with aberrantly activated myofibroblasts, forming a lesion vascularized by ectopic bronchial vessels in the presence of profibrotic monocyte-derived macrophage. The identification and detailed description of aberrant cell populations in the IPF lung may lead to identification of novel, cell-type specific therapies and biomarkers. Lastly, our lung cell atlas (*8*) provides an exhaustive reference of pulmonary cellular diversity in both healthy and diseased lung.

## Acknowledgments

We are indebted to all patients and control subjects who participated in this study. The sequencing was conducted by Mei Zhong at Yale Stem Cell Center Genomics Core facility. We thank Amos Brooks and the staff of Yale Pathology Tissue Services for tissue processing. We also thank Martijn C. Nawijn for providing access to his group’s data.

## Authors Contributions

N.K. and I.O.R conceptualized, acquired funding and supervised the study. B.R., G.R.W. performed sample collection and phenotyping. S.P., E.A, S.G.C. procured and dissociated the lungs. T.S.A., J.C.S., S.P., F.A., G.D. performed library construction. Data was processed, curated and visualized by T.S.A., J.C.S., N.N., X.Y. and analyzed by T.S.A, J.C.S., N.K., X.Y., N.N., M.J., Q.D., H.A., A.S. The online tool “IPFCellAtlas.com” was developed by N.N. Immunohistochemistry was performed by J.C.S. and T.S.A, and evaluated by R.H., N.K., J.C.S. The manuscript was drafted by T.S.A, J.C.S, N.K., and was reviewed and edited by all other authors.

## Data and materials availability

All raw count expression data was deposited to the Gene Expression Omnibus under accession number GSE136831. A user-friendly and accessible online tool for our data is available at www.IPFCellAtlas.com.

