## Supplemental_Materials for "Single Cell RNA-seq reveals ectopic and aberrant lung resident cell populations in Idiopathic Pulmonary Fibrosis"

**This PDF file includes:**

Materials and Methods
Supplementary Text
Supplementary References
Figs. S1 to S8
Tables S1 to S4
Captions for Data S1 to S9

**Other Supplementary Materials for this manuscript include the following:**

Data S1 to S9
[Data_S1_avgSubjectCelltype.markers.epiOnly.Final.txt,
Data_S2_avgSubjectCelltypeMarkers.mesenchymeOnly.Final.txt,
Data_S3_avgSubjectCelltypeMarkers.myeloidOnly.Final.txt,
Data_S4_avgSubjectCelltypeMarkers.lymphoidOnly.Final.txt,
Data_S5_AllCellTypesMarkers.logDOR.Final.txt,
Data_S6_CellCountsPerTypePerSubject_08152019.txt,
Data_S7_IPFvCtrl.wilcoxDE.collapsed.08292019.txt,
Data_S8_COPDvCtrl.wilcoxDE.collapsed.08292019.txt,
Data_S9_IPFvCOPD.wilcoxDE.collapsed.08292019.txt]

### Materials and Methods

Sample preparation for single cell sequencing

IPF and COPD lungs were obtained from patients undergoing transplant while healthy lungs were from rejected donor lung organs that underwent lung transplantation at the Brigham and Women's Hospital or donor organs provided by the National Disease Research Interchange (NDRI). The study protocol was approved by the Partners Healthcare Institutional Board Review (IRB Protocol # 2011P002419). Lungs were sliced and washed with cold sterile PBS. Visible airway structures, vessels, blood clots and mucin were removed. Lung specimens were minced mechanically into small pieces (<1mm$^3$) and then incubated for 45 minutes in 37°C in modified medium containing DMEM/F12K with added digestion enzymes (30 U/ml elastase (Elastin Products Company, Owensville, MO), 0.2 mg/ml DNAse I (Sigma, St. Louis, MO), 0.3 mg/ml liberase (Roche, Basel, Switzerland) and 1% Penicillin/Streptomycin). Digested tissue was filtered using a metal strainer. Unfiltered tissue was incubated a second time in digestion medium for 30 minutes, followed by repeat filtration. 10% FBS was added to the flow through to stop the enzymatic reaction. Product from filtration was centrifuged at 300G at 4°C for 10 minutes to collect cells dissociated from the tissue. Cell pellets were resuspended in Red Blood Cell Lysis Buffer (VWR International, Radnor, PA) for 3 minutes in 37°C and centrifuged again. The pellet was resuspended in DMEM/F12 medium and filtered using a 100µm strainer. Cells were resuspended in freezing medium (10%FBS and 10% DMSO in DMEM/F12), aliquoted and stored in liquid nitrogen.

Samples were thawed in a water bath at 37°C, filtered through a 100µm Cell Strainer (Fisher Scientific, USA), and rinsed with 20ml cold (4°C) PBS + 10% heat-inactivated FBS (both Life Technologies, USA). Cell suspensions were centrifuged at 300g, 5min, 4°C. Supernatant was discarded, cells were resuspended in 200µl Dead Cell removal microbead solution and incubated at 4°C for 15min. 2ml 1x Binding buffer was added, cell suspensions were passed through a 70µm cell strainer (Fisher Scientific, USA), and loaded onto a, with 500µl 1x binding buffer pre-washed, MS column (Dead Cell removal kit and MS columns: Miltenyi Biotec, USA). Cell suspensions passed through the MS columns, and then the columns were rinsed with 2ml 1x binding buffer. Cell suspensions were centrifuged at 300g, 5min, 4°C, the supernatant discarded, and the cells resuspended in 1ml PBS + 0.04 BSA (New England Biolabs, USA). Cell suspensions were passed through a final 40µm Cell Strainer (Fisher Scientific, USA). For cell concentrations and viability, cells were stained with Trypan blue, and then counted on a Countess Automated Cell Counter (Thermo Fisher, USA).

Single cell barcoding, library preparation, and sequencing

Single cells were barcoded using the 10x Chromium Single Cell platform, and cDNA libraries were prepared according to the manufacturer's protocol (Single Cell 3' Reagent Kits v2, 10x Genomics, USA). In brief, cell suspensions, reverse transcription master mix and partitioning oil were loaded on a single cell "A" chip, then run on the Chromium Controller. Reverse Transcription was performed within the droplets at 53°C for 45min. cDNA was amplified for a 12 cycles total on a BioRad C1000 Touch thermocycler. cDNA size selection was performed using SpriSelect beads (Beckman Coulter, USA) and a ratio of SpriSelect reagent volume to sample volume of 0.6. cDNA was analyzed on an Agilent Bioanalyzer High Sensitivity DNA chip for qualitative control purposes. cDNA was fragmented using the proprietary fragmentation enzyme blend for 5min at 32°C, followed by end repair and A-tailing at 65°C for 30min. cDNA were double-sided size selected using SpriSelect beads. Sequencing adaptors were ligated to the cDNA at 20°C for 15min. cDNA was amplified using a sample-specific index oligo as primer, followed by another round of double-sided size selection using SpriSelect beads. Final libraries were analyzed on an Agilent

Bioanalyzer High Sensitivity DNA chip for qualitative control purposes. cDNA libraries were sequenced on a HiSeq 4000 Illumina platform aiming for 150 million reads per library.

Data Processing, Computational Analyses

Fastq Generation and Read Trimming

Basecalls were converted to reads with the software Cell Ranger's (v2.2) implementation mkfastq. Multiple fastq files from the same library and strand were catenated to single files. Read2 files were subject to 2 passes of contaminant trimming with cutadapt (v1.17): first for the template switch oligo sequence (AAGCAGTGGTATCAACGCAGAGTACATGGG) anchored on the 5' end; secondly for poly(A) sequences on the 3' end. Following trimming, read pairs were removed if the read 2 was trimmed below 30bp.

Cell Barcode and Unique Molecular Identifier (UMI) Demultiplexing, Alignment and Transcript Counting

Subsequent read processing was conducted with the zUMIs pipeline (v2.0)(*1*). Paired reads were filtered if either the cell barcode or UMI sequence had more than 1 basepair with a phred < 20. Reads were aligned with STAR (v2.6.0c) (*2*) to the human genome reference GRCh38 release 91 from ensemble (*3*). Collapsed unique molecular identifiers (UMIs) with reads that span both exonic and intronic sequences were retained as both separate and combined gene expression assays. The top 10,000 cell barcodes ranked by reads were output from each library.

Filtering Valid Cell Barcodes and QC

Cell barcodes representative of quality cells were delineated from barcodes of apoptotic cells or background RNA based on a threshold of having at least 12% of transcripts arising from intron spanning - or unspliced - reads: indicative of nascent mRNA (supplemental figure 1). Cells with less than 1000 transcripts profiled or more than 20% of their transcriptome of mitochondrial origin were then removed.

Gene Name Conversion

Genes were originally output in ensemble gene ID format. To improve the interpretability of the variables without making compromises to sensitivity, we converted ensemble gene IDs to HGNC format using the R package BioMart (*4*) only when an exact one-to-one translation was available.

Data normalization and Cell Population Identification

UMIs from each cell barcode - irrespective of intron or exon coverage - were retained for all downstream analysis. Raw UMI counts were normalized with a scale factor of 10,000 UMIs per cell and subsequently natural log transformed with a pseudocount of 1. Aggregated data was subject to Louvain cluster analysis for cell type identification using the R package Seurat (version 2.3.1) (*5*). Recursive clustering analysis of subpopulations of pure immune (PTPRC+), epithelial (EPCAM) and the remaining mesenchymal populations were conducted to improve the granularity of our cell annotations. Multiplet cell populations were identified as having a transcriptomic signature that resembled the resulting combination of 2 or more disparate cell type signatures that already existed in the dataset. Cell barcodes flagged as multiplets were not included in downstream analyses.

Generation of Cell Type Markers

Cell-type marker lists were generated with two separate approaches. In the first approach, we sought to adjust for unbalanced representation of different cell-types across subjects, and reduce the zero-inflation in the data to generate a more reliable p-value from a differential expression test. We collapsed the gene expression matrix values to one where each column represents average gene expression for a given subject, for a given cell-type. In order to highlight gene expression among relatively similar cell-types, we then grouped each cell-type into one of four groups: epithelial, stromal/mesenchymal, myeloid and lymphoid. We then applied the Seurat FindAllMarkers implementation of the Wilcoxon rank-sum test each of these four subsets of cell types to generate separate marker lists for each (supplemental data S1-S4)

The second approach to generating cell-type markers uses a binary classifier system to assess the utility of detecting a given gene - irrespective of its intensity of expression - for classifying a cell. For each cell type in the data, we identified the genes whose expression was log fold change >= 0.3 greater than the other cells in the data. We then calculated the diagnostics odds ratio for each of these genes, where we binarize the expression values by treating any detection of a gene (normalized expression value > 0) as a positive value, and zero expression detection as negative. We included a pseudocount of 0.5 to avoid undefined values, as:

$$DOR = \frac{(TruePositives + 0.5)/(\text{FalsePositives} + 0.5)}{(\text{FalseNegatives} + 0.5)/(\text{TrueNegatives} + 0.5)}$$

Where *TruePositives* represents the number of cells within the group detected expressing the gene (value > 0), *FalsePositives* represents the number of cells outside of the group detected expressing the gene, *FalseNegatives* represents the number of cells within the group with no detected expression, and *TrueNegatives* represents the number of cells outside of the group with no detected expression of the gene. The log transformed DOR marker values are contained in supplemental table x.

UMAP Visualizations

For similarity-based cell network analysis and visualization, we utilized tools from the Python (version 3.6.8) library Scanpy (version 1.3.7) (*6*). Uniform Manifold Approximation and Projection (UMAP) (*7*) figures for all subsets of cells were generated using the same sequence of implementations. Feature selection of the top genes ranked by dispersion (scaled variance/mean) across 20 bins of the expression distribution are identified. The expression values for these genes are then adjusted for differences in total UMI and the fraction of mitochondrial reads across cells during z-normalization with a maximum absolute z-score of 10. The scaled values are then subject to principle component analysis (PCA) for linear dimension reduction.

The top principle components are subject to exploratory analysis to identify contributions to variance at the level of library batch, subject, cell type and disease. Additionally, the residuals of each principle component were explored to ascertain functional relevance of the signatures. Feature selection of principle components was conducted based on these analyses, and a shared nearest neighbor network was then created based on Euclidean distances between cells in multidimensional PC space and a fixed number of neighbors per cell, which was used to generate an intermediate 2-dimensional UMAP.

The resulting network was then subject to partition-based graph abstraction (PAGA) (*8*) using the cell type categories as abstraction nodes. A confidence threshold of cell type inter-connectivity was implemented to avoid spurious manifold connections between cell types of disparate lineages. Diffusion maps is then applied for non-linear dimension reduction on a more limited subset of principle component dimensions, and a new neighborhood graph is then computed based on representations of diffusion map distances. A final two-dimensional UMAP is then generated using the new neighborhood graph distances, initialized by the positions from the thresholded PAGA.

The values for all non-default parameters used during the generation of each UMAP are represented in supplemental table S3; if a parameter is unspecified, the default Scanpy implementation parameter was used.

Evaluation for the potential influence of outlying subject-specific variation

To ensure that our single cell analysis was not unduly driven by the outlying characteristics of one or few patient transcriptional profiles, we ran a parallel analysis using deep generative modeling to “correct” for any potentially aberrant subject-specific variational signatures in accordance with the single-cell Variational Inference (scVI) approach described by Lopez *et al*. (*9*). This was implemented through a dedicated python library downloaded from GitHub (https://github.com/YosefLab/scVI). We found that, even after applying this “batch effect” normalization, our cluster groups were preserved with high fidelity as evaluated by multiple metrics (Fig. S2), reinforcing that outlying subject-specific effects were not driving the results of our analysis.

Evaluation for the potential influence of cell cycle state

To further demonstrate the robustness of our approach, we applied a similar method to evaluate for possible data artifacts specific to cell cycle state. In another parallel analysis, we employed the approach described by Scialdone *et al*. (*10*), to extract G1, G2/M, and S components from each individual cell prior to normalization, using a set of 94 known cycle-associated genes. We then regressed out for these elements using Scanpy, similar to our approach for mitochondrial RNA, and proceeded with downstream analysis. We found that the G1, G2/M, and S states were well-distributed at both a cluster-level and patient-level. Furthermore, regressing out the effects of cell cycle state did not meaningfully alter our resulting cluster configurations (Fig. S3).

Comparison of manual cell annotations with automated methods

We compared our manual annotations to those produced through automated classification using SingleR (*11*). Strong correlation with manual annotations was found using both the HPCA and ENCODE human reference sets across our dataset, comparable to the correlation between the HPCA and ENCODE annotations themselves for these cells (Fig. S4).

PAGA Connectivity Analysis of Fibroblast and Myofibroblast

To assess the most likely trajectories of cell progression towards IPF-enriched fibrosis amongst fibroblast and myofibroblast, we used unsupervised Louvain (*12*) clustering to generate 8 subpopulations which were then subjected PAGA (*8*) analysis to ascertain the most likely inter-sub-cluster trajectories. The edge confidences between each subcluster node for all edges is visualized using the R package igraph (v1.2.4.1).

Scoring of regulon activity

A regulon is a group of target genes regulated by a common transcription factor. To score the activity of each regulon in each non-immune cell, we utilized the package pySCENIC (*13*) with default settings and the following database:

cisTarget databases (hg38__refseq-r80__500bp_up_and_100bp_down_tss.mc9nr.feather, hg38__refseq-r80__10kb_up_and_down_tss.mc9nr.feather) and the transcription factor motif annotation database (motifs-v9-nr.hgnc-m0.001-o0.0.tbl) were downloaded from resources.aertslab.org/cistarget/, the list of human transcription factors (hs_hgnc_tfs.txt) was downloaded from github.com/aertslab/pySCENIC/tree/master/resources.

Archetype Analysis of Fibroblast and Myofibroblast

We observed that many IPF-enriched features in the data were represented by a continuum of increasing phenotypic deviation from controls, rather than discrete features readily amenable to delineation (e.g. cluster analysis) from control-enriched signals. Consequently, we sought to implement archetype analysis of these continua to assess disease-enriched features rather than relying on traditional group-wise comparisons.

We first assessed the most likely trajectories towards fibrotic archetypes amongst fibroblast and myofibroblast using the diffusion pseudotime (DPT) (*14, 15*) implementation from Scanpy (*6*) to plot the distances along the UMAP manifold towards each archetype's terminus. The same diffusion map component structure used to generate the 2D UMAP visualizations were used for calculating DPT distances. For fibroblast, we took the cell with the highest expression of ITGB1, which lied at the terminus of fibroblast manifold's tendril, as the root cell and calculated the relative distances of all other fibroblast cells. We used 1-DPT value for the final distance value ordering and assigned a gradient of colors to cells for the range of 0 - 0.7 to represent each cells distance on the UMAP legend as well as in the accompanying heatmap.

For Myofibroblast, we calculated DPT ordering from all 3 termini in the myofibroblast manifold: using the highest MMP11 expressing cell to represent the IPF-enriched terminus, the furthest control cell to represent the control and COPD enriched terminus and the furthest cell from 225I to represent the single-subject-enriched terminus. To mitigate the impact on our analysis from the single-subject-enriched archetype, we removed the nearest 500 cells to the subject “225I” enriched terminus that did not overlap with the nearest 399 cells to the control and COPD enriched myofibroblast archetype. Amongst the remaining myofibroblast cells, we used the difference between both 1-DPT distance values from the remaining two non-subject-specific archetypes as a singular distance vector. A color gradient for the final 1-DPT range of -1 to 1 was then applied to each unfiltered cell in the UMAP legend and accompanying heatmap.

For both archetype heatmaps, cells were plotted in order of the final 1-DPT values from lowest to highest. A spearman correlation test was conducted between each cell's 1-DPT value and gene expression to ascertain which genes increased or decreased in expression along the manifold and would thus serve as candidate features in the heatmap. In addition to each cell's DPT distance value associated color, each cell's respective disease and subject color identity were included in the annotation bar to represent the extent of disease enrichment and subject-level contribution to the feature.

Archetype Analysis of Classical Monocytes and Macrophage

Amongst classical monocytes and Macrophage, we identified one monocyte archetype connected to distinct four macrophage archetypes: an inflammatory archetype, an IPF enriched archetype, an outlier MT-tRNA enriched archetype driven by two subjects and another outlier archetype driven by two separate subjects characterized by heatshock protein expression. 1-DPT distance values for all monocytes and macrophage were calculated from each archetype terminus as described for fibroblast and myofibroblast. We removed 1700 and 5800 most proximal cells to the MT-tRNA and heat shock protein archetypes respectively, to avoid contributions from outlier signals.

Following outlier removal, the 1-DPT distances values from the monocyte, inflamed macrophage and IPF-enriched macrophage were each independently unity normalized to values between 0 and 1. Distances along the three normalized trajectories were then used plot the cells in a ternary plot using the R gglot2 extension package tricolore (v1.2.0) to assign a color to each cell based on its relative proximity to each of the three archetype termini, which is visualized in both the UMAP color legend and ternary plot legend itself. The 15,000, 20,000 and 25,000, closest cells to the monocyte, inflamed macrophage and IPF-enriched macrophage archetype respectively were then selected for correlation analysis between 1-DPT values from each respective trajectory and gene expression to assess candidate genes for plotting the heatmap.

Gene Regulatory Network Construction

Gene regulatory networks (GRNs) for both control and IPF cell populations were generated using the r package bigSCale2 (*16*). Control and IPF cells were split apart and for each disease state, cells were randomly down-sampled using Seurat's SubsetData implementation (seed=7) to a maximum of 500 cells per cell-type, resulting in 10,560 cells from control and 15,388 cells from IPF. The resulting matrices were then filtered to remove genes with ensembl identifiers, and passed to bigSCale2, where both networks were constructed under the "normal" clustering parameter, with an edge cutoff of the top 0.993 quantile for correlation coefficient. Networks were visualized with the R package igraph, each network's layout is derived from 10,000 iterations of the Fuchterman-Reingold algorithm and "nogrid" parameter (seed=7).

Assessing Cell Relevance Scores to GRN Communities

Each GRN was clustered with igraph's "cluster_fast_greedy" function (seed=7).
Similar cell types were collapsed into the groups to avoid excessive phenotypic overlap: all dendritic cells; both macrophage varieties; both monocyte varieties; all T cells; NK & ILCs; both B cell varieties; both secretory; PNEC & ionocytes. For each cell-type group, we then took the top 500 genes ranked by log(DOR) markers as representative features. The relevance score for each cell-type group in a network's community is a z-score of the cumulative log(DOR) markers that intersect with members of each GRN community, weighted by the gene's normalized PageRank centrality within its community as follows:

$$\mathrm{PR}^*_{ij} = \frac{PR_{ij} - PR_{min.j}}{PR_{max.j} - PR_{min.j}}$$

$$\mathrm{CumulativeScore}_{kj} = \sum_{i \in kj} \log(\mathrm{DOR})_{ik} \times PR^*_{ij}$$

Where PR denotes the PageRank centrality for a node in the network and DOR is the diagnostic odds ratio classifier calculated earlier in the cell-type marker list. Given I genes, J communities and K cell-type groups, let $\mathrm{PR}^*_{ij}$ denote the normalized PageRank value for each gene i (i = 1, …, I) within its respective network community j (j = 1, …, J); let $\log(\mathrm{DOR})_{ik}$ be the natural log transformed DOR for gene i that belongs to cell-type group k (k = 1, …, K); let $i \in kj$ represent the intersection of genes from cell-type group k and GRN community j, and $\text{CumulativeScore}_{kj}$ represents the cumulative relevance score for cell-type group k in GRN community j. Lastly, the cumulative scores are then converted to z-scores by centering and scaling across cell-type groups within each community, only cell-type groups with a relevance score greater than or equal to 1 were retained for annotation of the community.

The aberrant basaloid cell-type community were excluded from the control GRN analysis, because this population of cells were not detected in any control samples.

Immunohistochemistry

The FFPE blocks were cut at 5 µm, then rehydrated using by standard xylene/ethanol deparaffinization. For Heat-induced antigen retrieval, specimens were boiled at 95°C for 20min in 1x Tris-Based Antigen Unmasking Solution (Vector Labs, USA). Tissue slides were incubated for 10min in BLOXALL Endogenous Peroxidase and Alkaline Phosphatase Blocking Solution (Vector Labs, USA) to block endogenous peroxidase and alkaline phosphatase activity. Tissue slides were blocked using 2.5% Normal Horse Serum Blocking Solution (Vector Labs, USA) for 20min, then incubated with the primary antibody (supplemental table S4), diluted in 2.5% Normal Horse Serum Blocking Solution, for 30min at room temperature. Specimen were incubated for 30min with secondary antibodies (anti-mouse or anti-rabbit ImmPRESS reagent, conjugated with horseradish peroxidase or alkaline phosphatase, as appropriate, all Vector Labs, USA). Slides were incubated for 10min in DAB working solution, or for 30min in Vector Reds working solution, as appropriate (both Vector Labs, USA). For sequential double-stainings, this protocol was repeated from the protein-blocking step on, using the alternative enzyme and substrate reagents. Tissue slides were counterstained in Hematoxilin Solution Gill #1 (Sigma Aldrich, USA) for 3min, then washed with tap water. For permanent mounting, specimens were dehydrated in Ethanol/Xylene, and then mounted VectaMount permanent mounting solution (Vector Labs, USA). Stained slides were digitalized on a Aperio Scanner (Leica), then analyzed using the softwares QuPath and ImageJ.

Independent validation of COL15A1 protein expression in peribronchial vascular endothelial cells

Images of COL15A1 immunohistochemical stainings of lung parenchyma and bronchi specimens were downloaded from the Human Protein Atlas ((*17*), https://www.proteinatlas.org/ENSG00000204291-COL15A1/tissue). All specimen had been stained using the polyclonal antibody HPA017915 (Sigma-Aldrich).

### Supplementary Text

Alternatively activated macrophage are a consistent feature of the IPF lung

Two groups conducting scRNAseq of IPF lungs have recently brought attention to a profibrotic macrophage found in IPF (*18*, *19*). To further elaborate on the features of these cells, we applied archetype analysis in a ternary fashion to classical monocytes, an IPF-specific macrophage archetype and a control-enriched inflammatory macrophage archetype (Fig. S8). We observed a sequential shift in features along the IPF macrophage archetype as it approached its most extreme terminus (Fig. S8D-E), SPP1 and cholesterol esterases LPL and LIPA expression increasing relatively early, ECM remodeling genes SPARC, GPC4, PALLD, CHI3L1, CTSK, MMP9 and MMP7 ramping up in expression further along the manifold. At the terminus of the IPF archetype macrophage start expressing CSF1, suggesting there's an autocrine feedback loop for recruitment and activation.

**Fig. S1.**

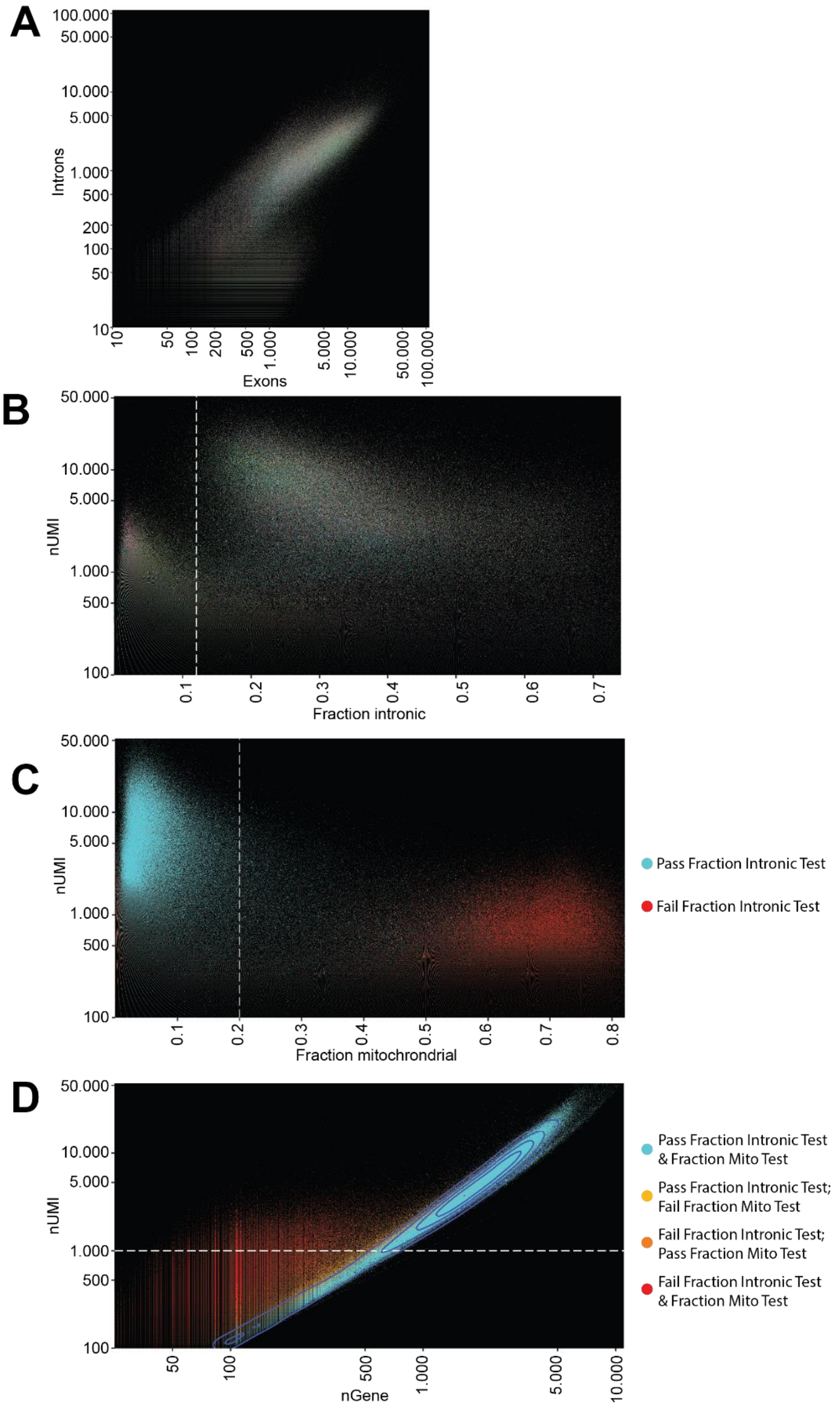


**(A)**. The relationship between the number intron-spanning unique molecular identifiers (UMIs) and exon-spanning UMIs, for the top 10,000 cell-barcodes per library ranked by number of UMIs, for all 107 libraries used in the data. The same cell-barcodes are plotted in all subsequent figures of the panel. **(B)** Relationship between the fraction of UMIs that are intron spanning and the total number of UMIs, the dotted line represents the cutoff of having at least 0.12 proportion intron-spanning to qualify as a valid cell. **(C)** The relationship between the fraction UMIs of mitochondrial origin and total number of UMIs. The dotted line represents the threshold of having no more than 0.2 proportion mitochondrial reads to qualify as a valid cell. cell-barcodes are colored based on whether they passed the previous test. **(D)** Relationship between the number of unique genes and the total number of UMIs for cell barcodes, blue contour lines represent population density estimation, the dotted line represents the cutoff of having at least 1000 unique genes to qualify as a valid cell. Cell barcodes are colored by whether they passed or failed previous thresholding tests.

**Fig. S2. Controlling for batch effects does not alter overall data architecture**

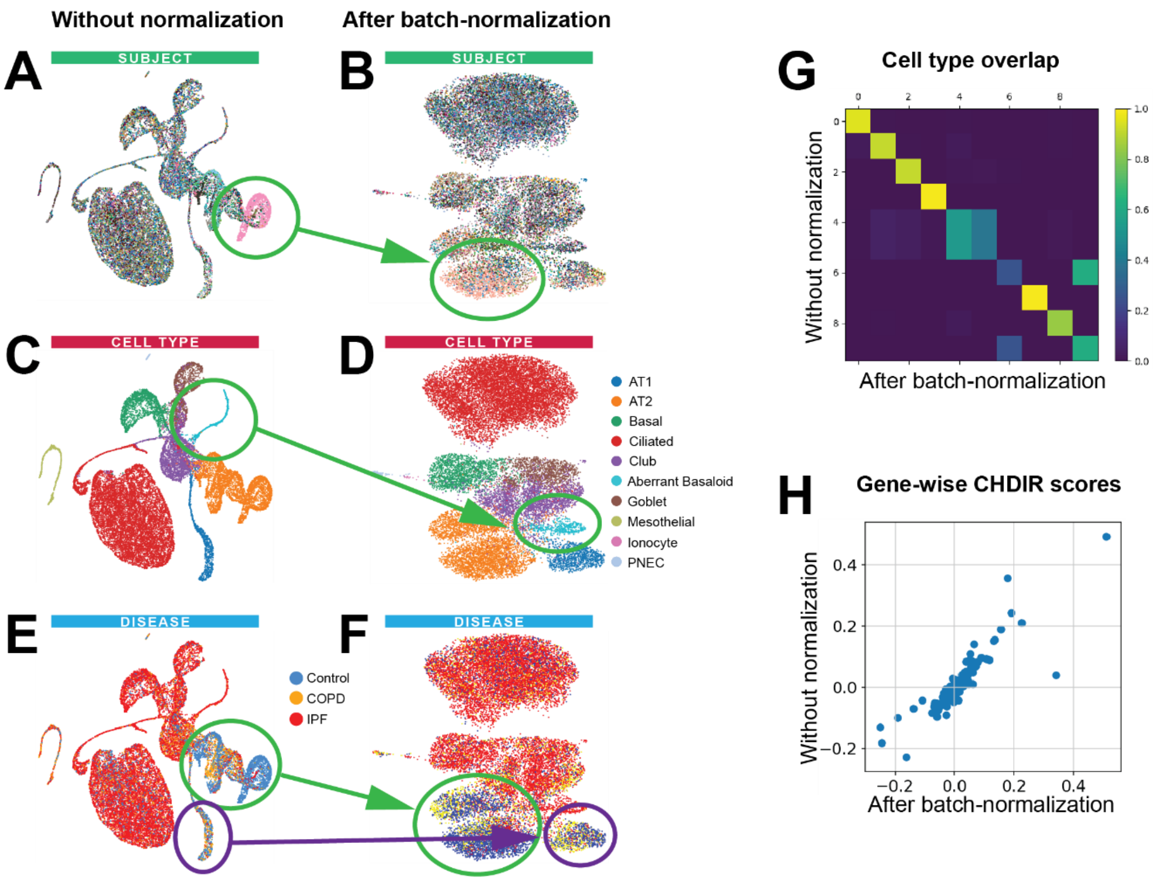


**I** **Without normalization**

| Index | Name | P-value | Adjusted p-value | Z-score | Combined score |
|---|---|---|---|---|---|
| 1 | Extracellular matrix organization | 1.08e-15 | 5.09e-13 | -2.11 | 72.73 |
| 2 | Integrin cell surface interactions | 1.20e-9 | 2.82e-7 | -2.00 | 41.00 |
| 3 | Assembly of collagen fibrils and other multimeric structures | 6.09e-8 | 8.27e-6 | -2.01 | 33.45 |
| 4 | Axon guidance | 1.10e-6 | 8.58e-5 | -2.32 | 31.85 |
| 5 | Non-integrin membrane-ECM interactions | 1.80e-7 | 1.69e-5 | -2.02 | 31.30 |
| 6 | ECM proteoglycans | 7.07e-8 | 8.27e-6 | -1.85 | 30.51 |
| 7 | Developmental Biology | 4.14e-6 | 2.42e-4 | -2.32 | 28.79 |
| 8 | Collagen formation | 2.16e-6 | 1.44e-4 | -1.97 | 25.74 |
| 9 | Hemostasis | 4.23e-5 | 1.80e-3 | -2.07 | 20.82 |
| 10 | Dissolution of Fibrin Clot | 6.46e-6 | 3.36e-4 | -1.71 | 20.45 |

**J** **After batch-normalization**

| Index | Name | P-value | Adjusted p-value | Z-score | Combined score |
|---|---|---|---|---|---|
| 1 | Extracellular matrix organization | 8.72e-12 | 3.97e-9 | -2.11 | 53.75 |
| 2 | Assembly of collagen fibrils and other multimeric structures | 3.01e-9 | 6.84e-7 | -2.03 | 39.74 |
| 3 | Collagen formation | 1.30e-8 | 1.97e-6 | -2.01 | 36.50 |
| 4 | Integrin cell surface interactions | 4.71e-6 | 5.36e-4 | -1.98 | 24.23 |
| 5 | Dissolution of Fibrin Clot | 6.46e-6 | 5.88e-4 | -1.87 | 22.30 |
| 6 | Degradation of the extracellular matrix | 1.13e-5 | 8.60e-4 | -1.94 | 22.10 |
| 7 | NCAM1 interactions | 3.19e-5 | 2.07e-3 | -1.97 | 20.36 |
| 8 | Collagen biosynthesis and modifying enzymes | 3.92e-5 | 2.23e-3 | -1.95 | 19.78 |
| 9 | Laminin interactions | 7.40e-5 | 3.60e-3 | -1.78 | 16.95 |
| 10 | Axon guidance | 7.07e-4 | 2.47e-2 | -2.26 | 16.39 |

**(A-B)** UMAPs of lung epithelial cells **(A)** without and **(B)** with correction for batch effects using single cell variational inference (scVI), colored by subject identity. Uncorrected clustering leads to a single focus of subject-driven cell focus, which is partially mitigated following batch correction. **(C-D)** UMAPs recolored by cell type **(C)** without and **(D)** with batch correction. The IPF-specific epithelial subpopulation of interest is similarly identified in both configurations (92.3% overlap). **(E-F)** UMAPs recolored by disease state **(E)** without and **(F)** with batch correction. Three clear foci of non-diseased cells are similarly present in both representations. **(G)** Overlap between cell type identifications derived with and without scVI batch normalization, showing high correlation between both analyses. **(H)** Scatter plot of characteristic direction (CHDIR) scores for top genes in IPF-specific epithelial subpopulation from both analyses. **(I-J)** Top 10 Reactome (*20*) pathways overrepresented among the 200 most differentially expressed genes in the IPF-specific epithelial subpopulations derived without **(I)** and with **(J)** batch normalization, with overlapping pathways highlighted in parallel colors. Taken together, these findings demonstrate that batch correction does not significantly alter the identification of this rare epithelial subpopulation, does not significantly change its underlying gene expression profile, and does not fundamentally alter the inferred functionality of these cells.

**Fig. S3. Controlling for cell cycle state does not alter overall data architecture**

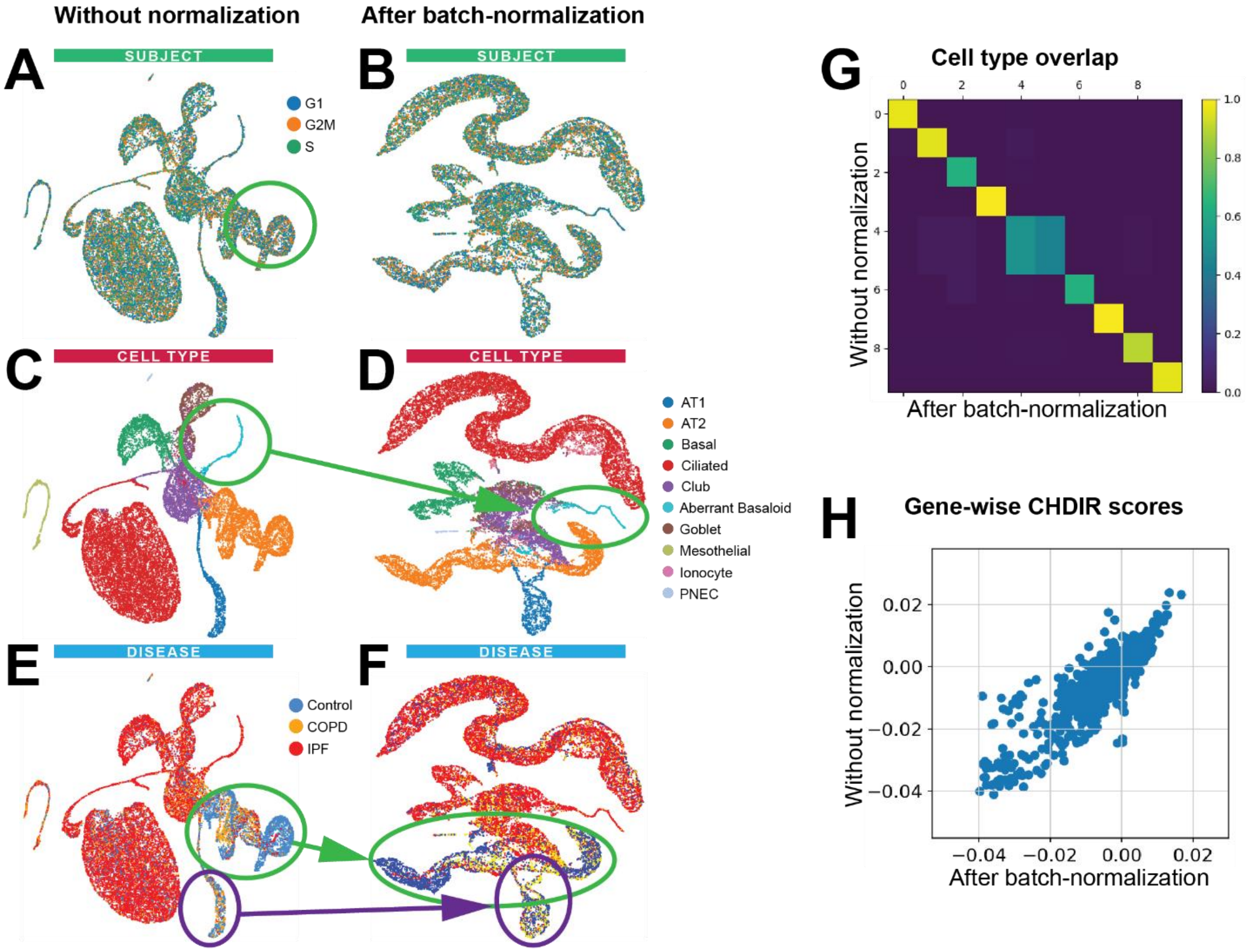


**I** **Without normalization**

| Index | Name | P-value | Adjusted p-value | Z-score | Combined score |
|---|---|---|---|---|---|
| 1 | Extracellular matrix organization | 1.08E-15 | 5.09E-13 | -2.11 | 72.73 |
| 2 | Integrin cell surface interactions | 1.20E-09 | 2.82E-07 | -2 | 41 |
| 3 | Assembly of collagen fibrils and other multimeric structures | 6.09E-08 | 8.27E-06 | -2.01 | 33.45 |
| 4 | Axon guidance | 1.10E-06 | 8.58E-05 | -2.32 | 31.85 |
| 5 | Non-integrin membrane-ECM interactions | 1.80E-07 | 1.69E-05 | -2.02 | 31.3 |
| 6 | ECM proteoglycans | 7.07E-08 | 8.27E-06 | -1.85 | 30.51 |
| 7 | Developmental Biology | 4.14E-06 | 2.42E-04 | -2.32 | 28.79 |
| 8 | Collagen formation | 2.16E-06 | 1.44E-04 | -1.97 | 25.74 |
| 9 | Hemostasis | 4.23E-05 | 1.80E-03 | -2.07 | 20.82 |
| 10 | Dissolution of Fibrin Clot | 6.46E-06 | 3.36E-04 | -1.71 | 20.45 |

**J** **After batch-normalization**

| Index | Name | P-value | Adjusted p-value | Z-score | Combined score |
|---|---|---|---|---|---|
| 1 | Extracellular matrix organization | 7.19E-11 | 2.45E-08 | -2.11 | 49.3 |
| 2 | Assembly of collagen fibrils and other multimeric structures | 1.07E-06 | 1.82E-04 | -2.03 | 27.84 |
| 3 | Collagen formation | 2.29E-05 | 2.10E-03 | -2.01 | 21.47 |
| 4 | Cell junction organization | 2.48E-05 | 2.10E-03 | -1.92 | 20.31 |
| 5 | Cell-Cell communication1 | 5.26E-05 | 3.57E-06 | -1.99 | 19.62 |
| 6 | Degradation of the extracellular matrix | 9.51E-05 | 4.62E-03 | -1.93 | 17.9 |
| 7 | Type I hemidesmosome assembly | 7.91E-05 | 4.48E-03 | -1.66 | 15.64 |
| 8 | Activation of Matrix Metalloproteinases | 2.80E-04 | 1.19E-02 | -1.9 | 15.52 |
| 9 | Cyclin D associated events in G1 | 5.48E-04 | 1.57E-02 | -2.07 | 15.51 |
| 10 | G1 Phase | 5.48E-04 | 1.57E-02 | -2.05 | 15.39 |

**(A-B)** UMAPs of lung epithelial cells **(A)** without and **(B)** with correction for cell cycle effects using single cell variational inference (scVI), colored by cell cycle phase. **(C-D)** UMAPs recolored by cell type **(C)** without and **(D)** with cycle correction. The IPF-specific epithelial subpopulation of interest is similarly identified in both configurations (89.4% overlap). **(E-F)** UMAPs recolored by disease state **(E)** without and **(F)** with cycle correction. Three clear foci of non-diseased cells are similarly present in both representations. **(G)** Overlap between cell type identifications derived with and without cell cycle normalization, showing high correlation between both analyses. **(H)** Scatter plot of characteristic direction (CHDIR) scores for top genes in IPF-specific epithelial subpopulation from both analyses. **(I-J)** Top 10 Reactome (*20*) pathways overrepresented among the 200 most differentially expressed genes in the IPF-specific epithelial subpopulations derived without **(I)** and with **(J)** batch normalization, with overlapping pathways highlighted in parallel colors. Taken together, these findings demonstrate that normalizing for cell cycle state does not significantly alter the identification of this rare epithelial subpopulation, does not significantly change its underlying gene expression profile, and does not fundamentally alter the inferred functionality of these cells.

**Fig. S4. Manual annotations of single cells are consistent with automated annotations drawn from multiple cell type definition databases.**

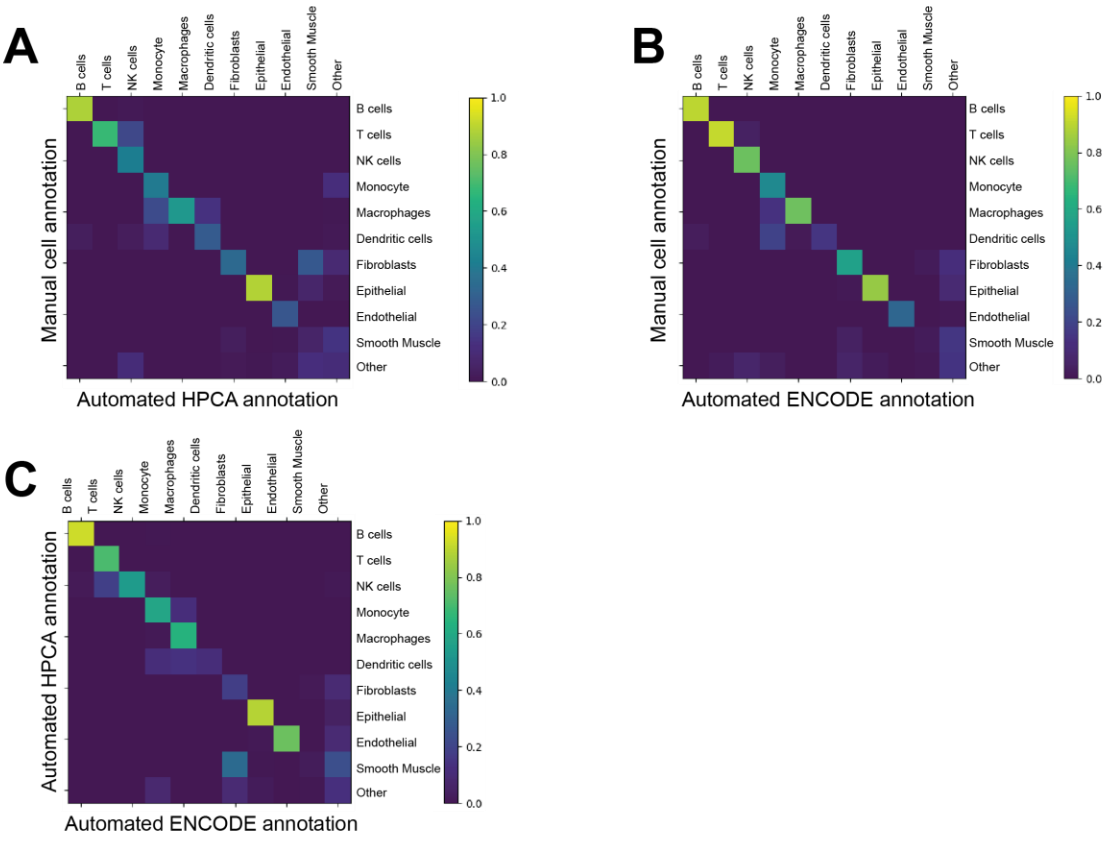


**D**

| HPCA Annotations | Aggregated Annotations |
|---|---|
| Astrocyte | Other |
| B_cell | B Cells |
| Chondrocytes | Other |
| DC | Dendritic Cells |
| Embryonic_stem cells | Other |
| Endothelial_cells | Endothelial Cells |
| Epithelial_cells | Epithelial Cells |
| Fibroblasts | Fibroblasts |
| GMP | Other |
| Gametocytes | Other |
| HSC_CD34+ | Other |
| MSC | Other |
| Macrophage | Macrophages |
| Monocyte | Monocytes |
| NK_cell | NK Cells |
| Neurons | Other |
| Neutrophil | Other |
| Neutrophils | Other |
| Osteoblasts | Other |
| Platelets | Other |
| Pre-B_cell_CD | Other |
| Pro-B_cell_CD34+ | Other |
| Smooth_muscle_cells | Smooth Muscle |
| T_cells | T Cells |
| Tissue_stem_cells | Other |
| iPS_cells | Other |

**E**

| Manual Cell Annotations | Aggregated Annotations |
|---|---|
| ATII_Epithelial_Cells | Epithelial Cells |
| ATI_Epithelial_Cells | Epithelial Cells |
| B | B Cells |
| B_Plasma | B Cells |
| Basal | Epithelial Cells |
| Ciliated | Epithelial Cells |
| Club | Epithelial Cells |
| Dendritic | Dendritic Cells |
| Dendritic_Langerhans | Dendritic Cells |
| Dendritic_Plasmacytoid | Dendritic Cells |
| Fibroblast | Fibroblast |
| Goblet | Epithelial Cells |
| Innate_lymphoid | Other |
| Ionocyte | Epithelial Cells |
| Lymphatic-Endothelial | Endothelial |
| Macrophage | Macrophage |
| Macrophage_Alveolar | Macrophage |
| Mast | Other |
| Mesothelial | Other |
| Monocyte | Monocytes |
| Myofibroblast | Fibroblast |
| Mystery Disease Epithelial | Epithelial Cells |
| NK | NK Cells |
| Neuroendocrine | Other |
| Pericytes | Other |
| Smooth-Muscle | Smooth Muscle |
| T | T Cells |
| T_Cytotoxic | T Cells |
| T_Regulatory | T Cells |
| Vascular-Endothelial | Endothelial |

**F**

| ENCODE Annotations | Aggregated Annotations |
|---|---|
| Adipocytes | Other |
| B-cells | B Cells |
| CD4+T-cells | T Cells |
| CD8+T-cells | T Cells |
| Chondrocytes | Other |
| DC | Dendritic Cells |
| Endothelial cells | Endothelial Cells |
| Epithelial cells | Epithelial Cells |
| Fibroblasts | Fibroblasts |
| HSC | Other |
| Keratinocytes | Other |
| Macrophages | Macrophages |
| Melanocytes | Other |
| Monocytes | Monocytes |
| Myocytes | Smooth Muscle |
| NK cells | NK Cells |
| Neurons | Other |
| Neutrophils | Other |
| Pericytes | Other |
| Smooth muscle | Smooth Muscle |

Using the SingleR software package, annotations were assigned to each cell in an automated fashion based on established definitions derived from either the **(A)** Human Primary Cell Atlas (HPCA) or **(B)** Blue ENCODE databases. Correlation scores were then assigned to pairwise comparisons of each definition based on the fraction of overlap between the two categories. (C) For reference, a comparison between the two automated databases is included, demonstrating comparable overlap. **(D-F)** Aggregation tables detailing the mappings of **(D)** HPCA, **(E)** Blue ENCODE, and **(F)** this manuscript's cell type definitions to the larger group categories used to derive scores in panels **(A-C)**.

**Fig. S5.**

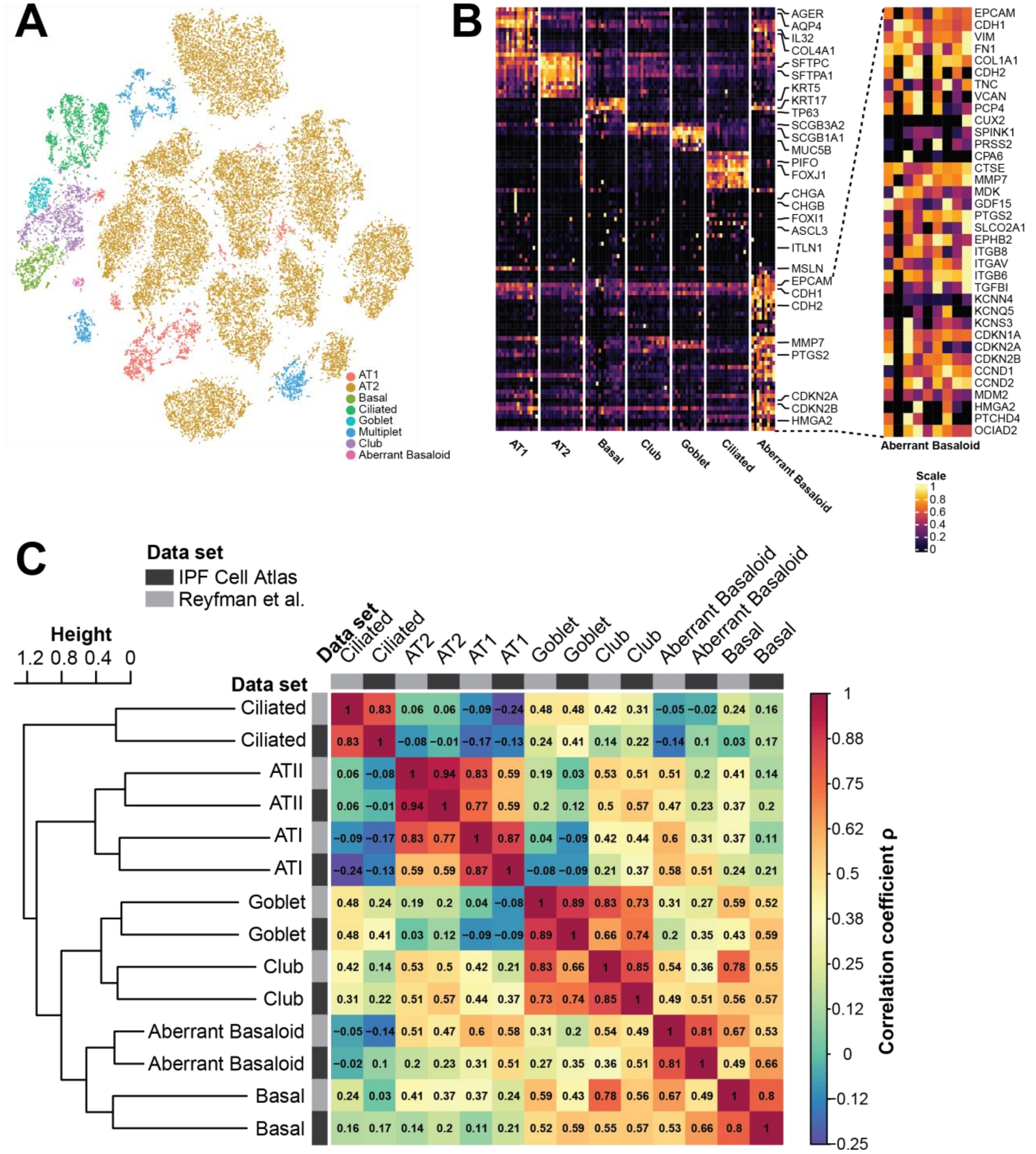


**(A)** UMAPs of all epithelial cells identified in Reyfman et al.'s (*18*) dataset labeled by cell type. **(B)** Heatmap of average gene expression per subject for each of the identified epithelial cell types in Reyfman et al.' s dataset. Column are hierarchically ordered by disease status and celltype. Right: Zoom annotation of distinguishing markers for aberrant basaloid cells identified in Reyfman et al.' s dataset. The portrayed genes and the order of the genes are the same as in figure 2C of the main manuscript. **(C)** Enlarged heatmap of the correlation matrix of figure 2E in the main manuscript, with the Spearman correlation coefficients ρ added as numbers. The heatmap portrays the correlation matrix of ATI, ATII, goblet, club, ciliated and hyperplastic basaloid cell average gene expression with analogous cell populations we identified in Reyfman et al.'s (*18*) dataset. Matrix cells are colored by Spearman's rho, cell populations are ordered with hierarchical clustering. The origin dataset for each cell population is denoted by different greyscale tones in the annotation bars.

**Fig. S6.**

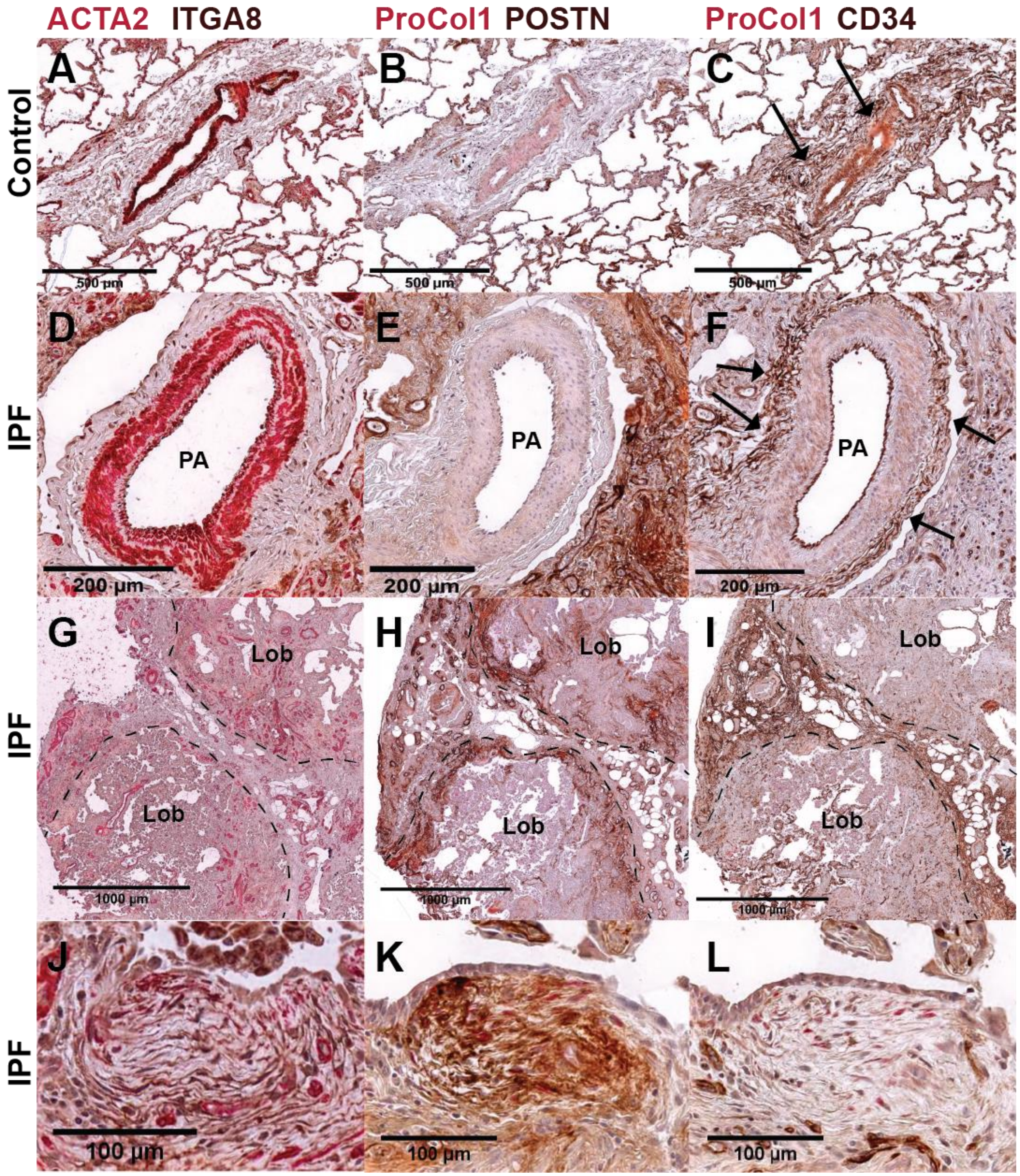


Immunohistochemistry of Control **(A-C)** and IPF **(D-L)** lung parenchyma. CD34+Fibroblasts are found in connective tissue surrounding pulmonary arteries **(C, F)** and in interlobular septae **(I, L)** in Control as well as IPF, while exhibiting no ITGA8 and POSTN staining. Myofibroblasts (ITGA8 and POSTN positive) are found in areas of dense fibrosis and especially in fibroblast foci **(J, K)**. Lob: pulmonary lobulus, PA: pulmonary arterial vessel, ProCol1: Pro-Collagen 1.

**Fig. S7. Validation of COL15A1 protein expression in peribronchial endothelial cells in the Human Protein Atlas**

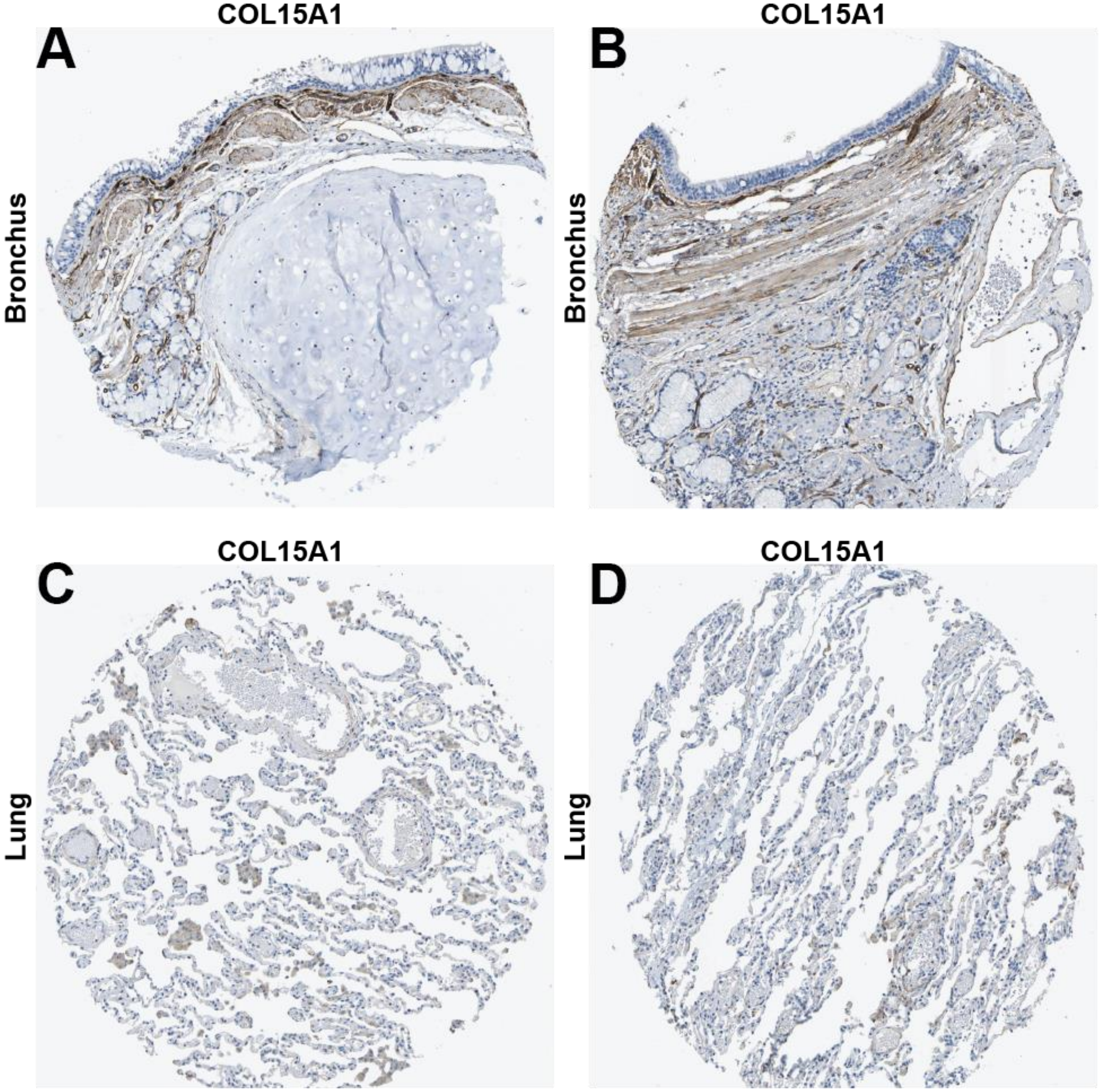


COL15A1 immunostainings - in brown - of bronchi **(A, B)** and lung parenchyma **(C, D)** specimen from the Human Protein Atlas (*17*). Pronounced COL15A1 positivity of peribronchial vessels **(A, B)**, while neither pulmonary capillaries nor larger pulmonary vessels stained positive for COL15A1 **(C, D)**.

**Fig. S8.**

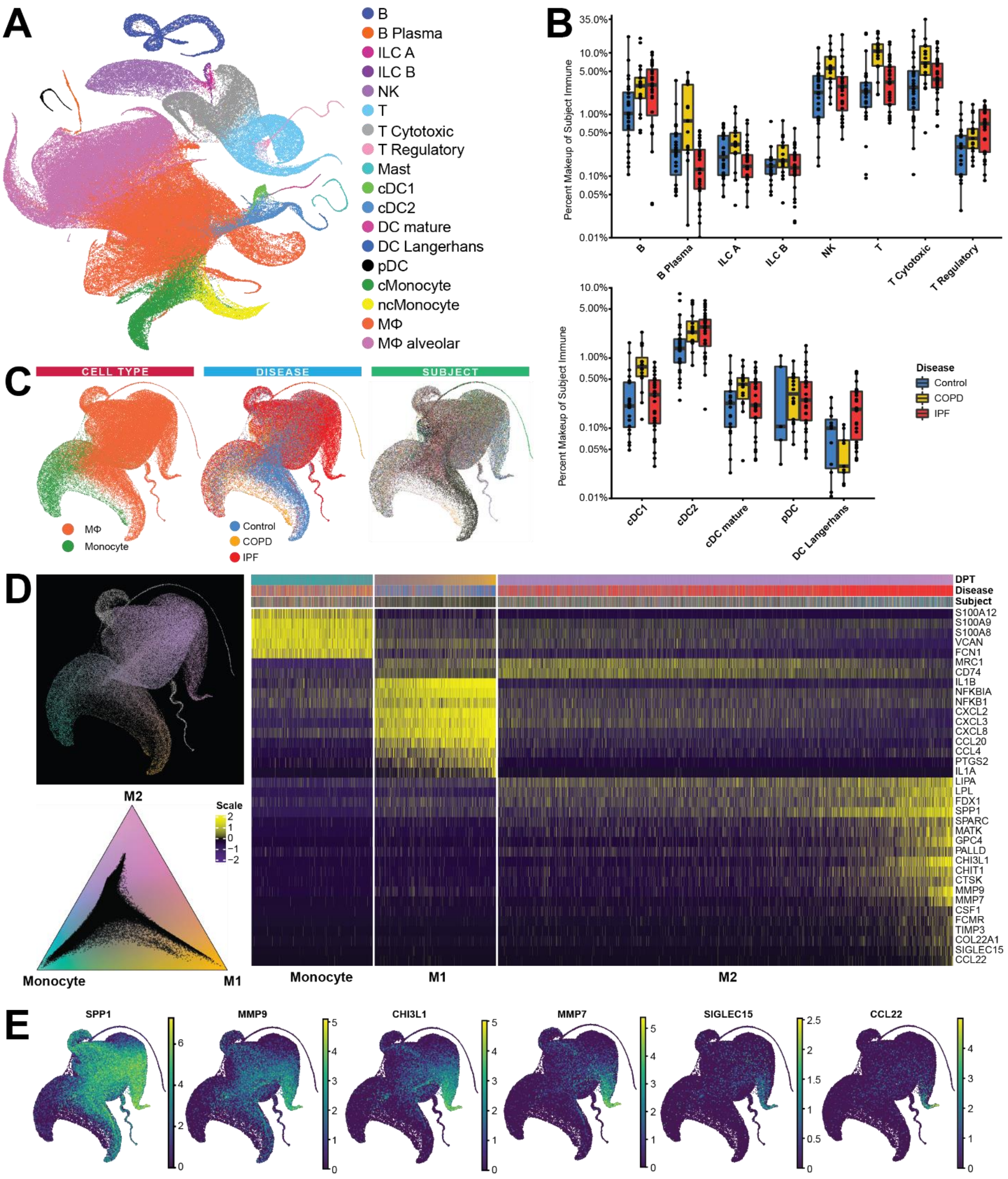


**(A)** UMAPs of immune cells, labelled by cell type, disease and subject. Each color represents a unique subject. **(B)** Boxplots representing the percent makeup distributions of each variety of immune cell as a proportion of all immune cells, within each disease group. Each dot represents a single subject, whiskers represent 1.5 x IQR. **(C)** UMAPs of classical monocytes and monocyte derived macrophages, labelled by cell type, disease and subject. Each color represents a unique subject. **(D)** Archetype analysis of classical monocytes and two macrophage UMAP manifold termini. Cells are colored with a ternary plot based on their global proximity to a monocyte terminus, an inflammatory macrophage terminus and an alternatively activated IPF enriched macrophage. Distance color assignments are also projected onto cells for UMAP visualization. The heatmap shows the most proximal cells to the monocyte terminus ordered from closest to furthest, while the most proximal cells to inflammatory macrophage and IPF enriched macrophage termini from furthest to closest. Each column represents a single cell, whose respective subject, disease, and DPT color assignment are in the annotation bar above. Macrophage cells belonging to two separate, single-subject enriched archetypes are removed from analysis. (E) UMAPs of monocyte and macrophage cells, colored by features that shift in the continuum towardsthe IPF-enriched macrophage archetype terminus, ordered by the distance in which the feature is detected, from furthest to nearest. Gene expression is normalized by natural log transformation with a pseudocount of 1, per 10,000 transcripts.

**Table S1.**

| Subject ID | Library IDs | Sex | Age | Race | Ever Smoker | Disease Group |
|---|---|---|---|---|---|---|
| 133C | 133C-a | Female | 32 | white | No | Control |
| 1372C | 137C-a, 137C-b | Female | 21 | white | No | Control |
| 034C | 034C | Male | 49 | asian | Yes | Control |
| 218C | 218C-a, 218C-b, 219C-a, 219C-b | Male | 29 | white | No | Control |
| 226C | 226C-a, 226C-b | Male | 32 | white | Yes | Control |
| 244C | 244C | Male | 50 | white | Yes | Control |
| 098C | 098C-a, 098C-b | Female | 41 | white | No | Control |
| 465C | 465C | Male | 56 | white | No | Control |
| 396C | 396C | Female | 37 | white | No | Control |
| 296C | 296C | Female | 80 | white | No | Control |
| 208C | 208C | Male | 23 | white | No | Control |
| 222C | 222C, 022C-a, 022C-b | Male | 65 | white | Yes | Control |
| 160C | 160C | Male | 64 | white | No | Control |
| 092C | 092C | Male | 29 | latino | No | Control |
| 439C | 439C, 439C-b | Female | 66 | white | No | Control |
| 065C | 065C | Female | 66 | white | No | Control |
| 388C | 388C | Male | 61 | white | No | Control |
| 192C | 92C, 192C-a | Female | 62 | white | No | Control |
| 483C | 483C | Male | 35 | white | No | Control |
| 001C | 001C | Male | 22 | white | No | Control |
| 002C | 002C | Female | 25 | white | No | Control |
| 003C | 003C | Female | 67 | white | No | Control |
| 454C | 454C | Female | 48 | white | No | Control |
| 253C | 253C | Female | 66 | white | Yes | Control |
| 484C | 484C | Male | 31 | white | Yes | Control |
| 081C | 081C | Male | 20 | white | No | Control |
| 137C | 137C | Male | 54 | white | No | Control |
| 084C | 084C | Male | 46 | black | No | Control |
| 152CO | 152CO, 152CO-a | Male | 57 | white | Yes | COPD |
| 153CO | 153CO-a, 153CO-b | Male | 62 | white | Yes | COPD |
| 178CO | 178CO | Female | 58 | white | Yes | COPD |
| 184CO | 184CO-a, 184CO-b | Female | 55 | white | No | COPD |
| 186CO | 186CO-b | Male | 66 | white | Yes | COPD |
| 192CO | 192CO, 192CO-a, 192CO-b | Male | 66 | white | Yes | COPD |
| 193CO | 193CO-a, 193CO-b | Male | 63 | white | Yes | COPD |
| 194CO | 194CO-a | Male | 59 | white | Yes | COPD |
| 207CO | 207CO | Female | 60 | white | Yes | COPD |
| 217CO | 217CO-a | Female | 70 | white | Yes | COPD |
| 23CO | 23CO | Male | 66 | white | Yes | COPD |
| 235CO | 235CO | Female | 61 | white | Yes | COPD |

| | | | | | | |
|---|---|---|---|---|---|---|
| 237CO | 237CO | Female | 57 | white | Yes | COPD |
| 238CO | 238CO | Male | 66 | white | Yes | COPD |
| 8CO | 8CO | Male | 65 | white | Yes | COPD |
| 052CO | 052CO-a | Female | 62 | white | Yes | COPD |
| 056CO | 056CO | Female | 57 | white | Yes | COPD |
| 137CO | 137CO | Female | 73 | white | Yes | COPD |
| 210I | 210CO | Male | 68 | white | Yes | IPF |
| 135I | 135I-a, 135I-b | Male | 59 | white | Yes | IPF |
| 138I | 138I, 138I-a | Male | 56 | white | Yes | IPF |
| 145I | 145I-a | Male | 67 | white | Yes | IPF |
| 157I | 157I, 157I-a, 157I-b | Female | 66 | white | Yes | IPF |
| 166I | 166I-a | Male | 63 | white | Yes | IPF |
| 174I | 174I-a | Female | 67 | white | Yes | IPF |
| 179I | 179I | Male | 70 | white | Yes | IPF |
| 209I | 209I-a | Female | 65 | asian | No | IPF |
| 212I | 212I-a | Male | 71 | white | No | IPF |
| 214I | 214I-a | Male | 69 | white | Yes | IPF |
| 221I | 221I-a | Female | 67 | white | No | IPF |
| 222I | 222I-a, 222I-b | Male | 59 | white | No | IPF |
| 225I | 225I-a, 225I-b | Female | 70 | white | Yes | IPF |
| 228I | 228I-a, 228I-b | Male | 56 | white | No | IPF |
| 29I | 29I | Male | 61 | white | No | IPF |
| 051I | 051I, 051I-a | Male | 62 | other | Yes | IPF |
| 025I | 025I | Male | 65 | white | Yes | IPF |
| 010I | 010I | Male | 78 | white | Yes | IPF |
| 021I | 021I | Male | 69 | white | No | IPF |
| 022I | 022I | Male | 67 | white | Yes | IPF |
| 041I | 041I-b | Male | 59 | white | No | IPF |
| 040I | 040I, 040I-b | Male | 70 | white | Yes | IPF |
| 47I | 47I-a, 47I-b | Male | 64 | white | No | IPF |
| 49I | 49I-a, 49I-b | Male | 66 | white | Yes | IPF |
| 053I | 53I, 053I-d, 053I-n | Male | 60 | white | Yes | IPF |
| 59I | 59I | Female | 66 | white | No | IPF |
| 063I | 063I-b | Male | 68 | white | No | IPF |
| 034I | 034I-a | Male | 54 | white | Yes | IPF |
| 123I | 123I | Male | 74 | white | Yes | IPF |
| 158I | 158I-b | Male | 68 | white | Yes | IPF |
| 177I | 177I | Male | 69 | white | Yes | IPF |

Basic characteristics of all included patients and controls. Age is given in years.

**Table S2.**

| Subject ID | Library ID | Raw Reads [*10^6] | TSO Trimmed [%] | PolyA Trimmed [%] | Pass Trimming [%] | Reads Pass Trimming [*10^6] | Pass Filter [%] | Reads Pass Filter [*10^6] | Mapped Unique [%] | Mapped Multi [%] | too short [%] | Splice Junctions [*10^6] | Non Canonical Splices [%] |
|---|---|---|---|---|---|---|---|---|---|---|---|---|---|
| 001C | 001C | 107.2 | 8.2 | 5.6 | 99.5 | 106.6 | 86.1 | 91.8 | 89.1 | 7.9 | 2.9 | 12.4 | 0.8 |
| 002C | 002C | 108.6 | 9.4 | 5.3 | 99.6 | 108.2 | 86.1 | 93.2 | 88.6 | 8.3 | 3.0 | 11.4 | 0.9 |
| 003C | 003C | 185.7 | 6.5 | 5.9 | 99.4 | 184.7 | 87.3 | 161.2 | 89.9 | 7.2 | 2.7 | 24.4 | 0.5 |
| 010I | 010I | 182.8 | 7.4 | 27.3 | 99.2 | 181.4 | 89.9 | 163.2 | 88.5 | 7.5 | 4.0 | 21.6 | 0.5 |
| 021I | 021I | 175.4 | 7.1 | 28.3 | 98.9 | 173.5 | 89.9 | 156 | 87.4 | 8.3 | 4.3 | 19.3 | 0.5 |
| 222C | 022C-a | 193.5 | 25.0 | 4.8 | 99.5 | 192.5 | 91.5 | 176.2 | 89.7 | 7.6 | 2.6 | 25.8 | 0.7 |
| 222C | 022C-b | 186 | 33.3 | 5.2 | 99.1 | 184.2 | 93.6 | 172.4 | 90.6 | 7.4 | 1.8 | 26.5 | 0.6 |
| 022I | 022I | 180.4 | 7.2 | 27.0 | 99.4 | 179.3 | 89.9 | 161.2 | 88.8 | 7.4 | 3.8 | 21.6 | 0.5 |
| 025I | 025I | 182.4 | 6.9 | 29.6 | 97.9 | 178.6 | 89.0 | 158.9 | 87.6 | 9.0 | 3.3 | 15 | 0.7 |
| 034C | 034C | 195.3 | 24.2 | 27.3 | 81.0 | 158.2 | 78.2 | 123.7 | 84.1 | 9.8 | 5.9 | 16.3 | 0.6 |
| 034I | 034I-a | 207 | 4.8 | 4.4 | 99.7 | 206.4 | 92.0 | 189.9 | 88.8 | 6.9 | 4.2 | 28.6 | 0.6 |
| 040I | 040I | 185.9 | 6.7 | 9.3 | 97.2 | 180.7 | 74.3 | 134.2 | 86.9 | 7.6 | 5.2 | 20.7 | 0.5 |
| 040I | 040I-b | 187.3 | 8.6 | 4.8 | 99.4 | 186.2 | 92.7 | 172.5 | 89.7 | 6.3 | 3.9 | 22.2 | 0.7 |
| 041I | 041I-b | 209 | 6.9 | 5.1 | 99.5 | 207.9 | 93.1 | 193.4 | 90.9 | 6.9 | 2.1 | 29.9 | 0.5 |
| 051I | 051I | 186.6 | 23.8 | 13.7 | 92.6 | 172.8 | 71.1 | 122.9 | 83.8 | 10.7 | 5.0 | 12.9 | 1.8 |
| 051I | 051I-a | 221 | 25.4 | 5.3 | 99.1 | 219 | 92.1 | 201.6 | 86.8 | 9.6 | 3.5 | 24.6 | 0.6 |
| 052CO | 052CO-a | 204.8 | 7.3 | 5.7 | 99.5 | 203.7 | 91.6 | 186.6 | 89.3 | 7.6 | 3.0 | 30.2 | 0.8 |
| 053I | 053I-d | 132.6 | 34.0 | 13.7 | 92.5 | 122.6 | 83.2 | 102 | 86.2 | 9.5 | 3.1 | 11.6 | 0.8 |
| 053I | 053I-n | 121.4 | 40.5 | 22.0 | 85.0 | 103.2 | 83.3 | 86 | 82.6 | 9.7 | 6.5 | 10.8 | 0.7 |
| 056CO | 056CO | 183.4 | 16.0 | 9.6 | 96.6 | 177.2 | 90.3 | 160 | 89.1 | 8.5 | 2.2 | 15.9 | 1.6 |
| 063I | 063I-b | 215.2 | 9.0 | 5.6 | 99.3 | 213.8 | 94.1 | 201.1 | 91.3 | 6.8 | 1.8 | 35 | 0.5 |
| 065C | 065C | 115.8 | 11.7 | 10.7 | 95.4 | 110.4 | 89.0 | 98.2 | 90.2 | 7.2 | 2.4 | 8.9 | 1.6 |
| 081C | 081C | 126.4 | 13.9 | 6.9 | 98.0 | 123.9 | 89.3 | 110.6 | 87.1 | 10.1 | 2.7 | 9.8 | 1.3 |
| 084C | 084C | 134.4 | 32.7 | 24.5 | 80.9 | 108.7 | 90.3 | 98.2 | 81.4 | 15.6 | 2.8 | 5.4 | 2.1 |
| 092C | 092C | 120.5 | 5.6 | 6.2 | 99.4 | 119.8 | 86.7 | 103.8 | 87.8 | 8.4 | 3.7 | 19 | 0.3 |
| 098C | 098C-a | 235.6 | 7.7 | 5.0 | 99.6 | 234.6 | 93.8 | 219.9 | 90.3 | 7.1 | 2.5 | 37.7 | 0.5 |
| 098C | 098C-b | 209.1 | 6.9 | 5.6 | 99.4 | 208 | 92.9 | 193.2 | 90.9 | 6.9 | 2.1 | 29.9 | 0.6 |
| 123I | 123I | 177.6 | 7.8 | 8.3 | 97.7 | 173.6 | 73.7 | 128 | 88.0 | 7.9 | 4.0 | 24.4 | 0.3 |
| 133C | 133C-a | 241.4 | 10.0 | 7.0 | 99.2 | 239.4 | 91.9 | 220.1 | 89.8 | 7.6 | 2.4 | 38.8 | 0.6 |
| 135I | 135I-a | 197 | 11.6 | 6.4 | 98.9 | 194.8 | 91.8 | 178.8 | 88.8 | 7.6 | 3.4 | 25.4 | 0.8 |
| 135I | 135I-b | 184.1 | 11.7 | 5.5 | 99.2 | 182.7 | 92.5 | 169 | 89.9 | 7.4 | 2.5 | 26.3 | 0.8 |
| 137C | 137C | 217.8 | 18.3 | 17.4 | 89.6 | 195.1 | 73.8 | 144 | 85.0 | 8.9 | 6.0 | 18.6 | 0.6 |
| 1372C | 137C-a | 179.3 | 9.6 | 6.9 | 98.6 | 176.9 | 91.8 | 162.5 | 89.6 | 7.9 | 2.3 | 22.7 | 0.8 |
| 1372C | 137C-b | 189.9 | 9.1 | 6.2 | 99.1 | 188.3 | 91.4 | 172.1 | 89.6 | 8.1 | 2.1 | 28.1 | 0.7 |
| 137CO | 137CO | 268.2 | 9.4 | 15.3 | 91.7 | 246.1 | 73.9 | 181.8 | 85.2 | 10.0 | 4.7 | 19.3 | 0.5 |
| 138I | 138I | 258.4 | 3.6 | 5.0 | 99.6 | 257.3 | 79.1 | 203.7 | 89.8 | 7.4 | 2.5 | 32.9 | 0.5 |
| 138I | 138I-a | 226.1 | 50.9 | 8.2 | 96.2 | 217.6 | 90.1 | 196.1 | 81.2 | 8.0 | 10.6 | 30.1 | 0.6 |

| | | | | | | | | | | | | | |
|---|---|---|---|---|---|---|---|---|---|---|---|---|---|
| 145I | 145I-a | 239.6 | 8.6 | 5.2 | 99.5 | 238.3 | 92.7 | 220.8 | 89.6 | 7.6 | 2.7 | 37.7 | 0.5 |
| 152CO | 152CO | 312 | 2.8 | 4.8 | 99.5 | 310.6 | 78.9 | 245 | 87.1 | 9.5 | 3.1 | 28.2 | 0.5 |
| 152CO | 152CO-a | 162.1 | 57.9 | 7.5 | 97.4 | 157.9 | 91.6 | 144.7 | 87.7 | 7.4 | 4.7 | 17.7 | 0.9 |
| 153CO | 153CO-a | 148 | 48.3 | 10.5 | 93.8 | 138.8 | 91.6 | 127.1 | 88.1 | 8.3 | 3.4 | 13.2 | 1.0 |
| 153CO | 153CO-b | 191.8 | 31.3 | 9.3 | 95.5 | 183.3 | 91.3 | 167.4 | 88.9 | 8.8 | 2.1 | 17.2 | 1.1 |
| 157I | 157I | 252.9 | 8.0 | 9.9 | 96.1 | 243.1 | 74.1 | 180.1 | 86.9 | 9.2 | 3.7 | 26.6 | 0.4 |
| 157I | 157I-a | 205.2 | 8.3 | 6.5 | 98.7 | 202.7 | 92.9 | 188.2 | 89.5 | 7.9 | 2.4 | 30.2 | 0.6 |
| 157I | 157I-b | 185.1 | 8.6 | 5.6 | 99.2 | 183.7 | 91.7 | 168.3 | 89.6 | 8.1 | 2.1 | 27.6 | 0.6 |
| 158I | 158I-b | 204.6 | 9.2 | 4.9 | 99.6 | 203.7 | 93.7 | 190.9 | 91.6 | 6.7 | 1.6 | 36 | 0.5 |
| 160C | 160C | 137.7 | 8.2 | 6.1 | 99.5 | 137 | 87.2 | 119.5 | 89.3 | 7.3 | 3.2 | 14.7 | 0.7 |
| 166I | 166I-a | 192.7 | 61.3 | 5.4 | 98.8 | 190.3 | 91.8 | 174.6 | 86.7 | 7.5 | 5.6 | 24.6 | 0.8 |
| 174I | 174I-a | 208.8 | 8.0 | 4.4 | 99.7 | 208.1 | 93.8 | 195.2 | 89.9 | 7.5 | 2.5 | 38.5 | 0.4 |
| 177I | 177I | 156.9 | 8.8 | 5.1 | 99.5 | 156.2 | 86.9 | 135.8 | 88.9 | 8.2 | 2.8 | 25.3 | 0.4 |
| 178CO | 178CO | 194.7 | 9.1 | 7.5 | 98.8 | 192.3 | 90.5 | 174 | 89.1 | 8.0 | 2.7 | 24.3 | 1.2 |
| 179I | 179I | 161.7 | 8.3 | 7.5 | 98.6 | 159.4 | 87.2 | 139 | 89.2 | 8.0 | 2.7 | 21.3 | 0.6 |
| 184CO | 184CO-a | 158.9 | 57.0 | 8.0 | 96.9 | 153.9 | 91.3 | 140.5 | 88.5 | 7.3 | 4.0 | 16.6 | 0.9 |
| 184CO | 184CO-b | 178.9 | 30.0 | 5.5 | 99.1 | 177.3 | 92.0 | 163.1 | 90.0 | 7.3 | 2.4 | 21.2 | 0.9 |
| 186CO | 186CO-b | 192.1 | 10.8 | 5.8 | 99.2 | 190.5 | 94.0 | 178.9 | 90.5 | 7.8 | 1.6 | 23 | 0.7 |
| 192C | 192C-a | 203.9 | 34.0 | 8.0 | 97.3 | 198.3 | 91.4 | 181.3 | 88.5 | 7.9 | 3.3 | 23.1 | 0.7 |
| 192CO | 192CO | 249.5 | 6.9 | 9.1 | 97.2 | 242.5 | 74.6 | 180.9 | 87.6 | 8.7 | 3.5 | 32 | 0.3 |
| 192CO | 192CO-a | 196.3 | 6.9 | 5.7 | 99.4 | 195.1 | 93.0 | 181.4 | 89.8 | 7.9 | 2.1 | 28.4 | 0.5 |
| 192CO | 192CO-b | 186.4 | 5.9 | 4.7 | 99.7 | 185.9 | 92.8 | 172.4 | 90.4 | 7.6 | 1.9 | 28.1 | 0.5 |
| 193CO | 193CO-a | 197.2 | 5.8 | 4.5 | 99.7 | 196.6 | 92.7 | 182.2 | 89.0 | 7.4 | 3.5 | 29.4 | 0.4 |
| 193CO | 193CO-b | 169.7 | 5.3 | 5.1 | 99.6 | 169 | 92.2 | 155.7 | 90.8 | 7.2 | 1.9 | 24.5 | 0.5 |
| 194CO | 194CO-a | 179.8 | 69.1 | 11.3 | 93.9 | 168.8 | 90.8 | 153.2 | 81.1 | 10.3 | 8.3 | 14.7 | 1.1 |
| 207CO | 207CO | 169.7 | 13.9 | 8.8 | 96.9 | 164.5 | 89.0 | 146.4 | 88.8 | 8.0 | 3.0 | 17.6 | 1.5 |
| 208C | 208C | 112.7 | 10.0 | 29.8 | 97.6 | 110 | 90.8 | 99.9 | 86.6 | 8.4 | 5.0 | 10.7 | 0.6 |
| 209I | 209I-a | 183.3 | 9.1 | 5.2 | 99.5 | 182.3 | 91.7 | 167.1 | 90.2 | 6.7 | 3.0 | 25.7 | 0.5 |
| 210I | 210CO | 227.5 | 10.5 | 6.3 | 98.4 | 223.9 | 88.4 | 198 | 88.9 | 8.2 | 2.8 | 20.4 | 0.8 |
| 212I | 212I-a | 166 | 16.4 | 6.7 | 98.8 | 164 | 91.8 | 150.6 | 88.8 | 7.8 | 3.2 | 18.1 | 0.8 |
| 214I | 214I-a | 170.8 | 58.1 | 14.1 | 90.4 | 154.4 | 91.3 | 141.1 | 87.0 | 8.8 | 4.0 | 10.1 | 1.5 |
| 217CO | 217CO-a | 185.2 | 24.5 | 5.4 | 99.2 | 183.6 | 91.5 | 167.9 | 88.3 | 7.4 | 4.2 | 21.2 | 0.8 |
| 218C | 218C-a | 162.2 | 48.2 | 16.3 | 88.9 | 144.1 | 91.4 | 131.7 | 86.5 | 8.6 | 4.8 | 17 | 0.9 |
| 218C | 218C-b | 197.5 | 36.7 | 7.6 | 96.7 | 191 | 91.6 | 175 | 88.5 | 8.5 | 2.8 | 22.2 | 0.9 |
| 218C | 219C-a | 149.9 | 67.3 | 8.4 | 95.7 | 143.5 | 91.4 | 131.1 | 87.2 | 7.8 | 4.8 | 17 | 0.8 |
| 218C | 219C-b | 196.8 | 35.0 | 9.8 | 94.9 | 186.8 | 91.7 | 171.3 | 89.4 | 7.9 | 2.5 | 25.1 | 0.8 |
| 221I | 221I-a | 167 | 13.9 | 5.4 | 99.3 | 165.8 | 91.7 | 152 | 88.8 | 7.3 | 3.7 | 23.3 | 0.5 |
| 222C | 222C | 180.4 | 7.4 | 5.3 | 99.6 | 179.6 | 87.1 | 156.4 | 89.5 | 7.6 | 2.7 | 19.5 | 0.7 |
| 222I | 222I-a | 167.3 | 8.0 | 4.8 | 99.6 | 166.6 | 91.6 | 152.5 | 90.0 | 6.9 | 3.0 | 25.9 | 0.5 |
| 222I | 222I-b | 198.7 | 8.8 | 5.4 | 99.4 | 197.6 | 91.6 | 181 | 90.3 | 7.3 | 2.2 | 29.1 | 0.6 |
| 225I | 225I-a | 198.1 | 9.6 | 7.2 | 99.1 | 196.3 | 91.6 | 179.9 | 90.2 | 7.0 | 2.6 | 23.8 | 0.8 |

| | | | | | | | | | | | | | |
|---|---|---|---|---|---|---|---|---|---|---|---|---|---|
| 225I | 225I-b | 209.4 | 7.7 | 4.4 | 99.7 | 208.9 | 92.3 | 192.8 | 91.2 | 6.4 | 2.2 | 28.6 | 0.7 |
| 226C | 226C-a | 179.7 | 60.0 | 8.0 | 97.4 | 175 | 91.9 | 160.9 | 87.8 | 7.5 | 4.6 | 19.1 | 0.8 |
| 226C | 226C-b | 238.1 | 35.4 | 5.6 | 99.0 | 235.7 | 93.0 | 219 | 89.8 | 7.6 | 2.4 | 35.1 | 0.8 |
| 228I | 228I-a | 218.9 | 5.7 | 4.2 | 99.7 | 218.3 | 93.1 | 203.2 | 90.9 | 6.3 | 2.7 | 18.3 | 1.1 |
| 228I | 228I-b | 217.4 | 4.5 | 5.3 | 99.6 | 216.6 | 92.2 | 199.7 | 90.2 | 7.0 | 2.7 | 22.2 | 1.0 |
| 235CO | 235CO | 111.6 | 16.5 | 8.4 | 97.3 | 108.6 | 88.5 | 96.1 | 88.8 | 7.7 | 3.3 | 11.8 | 0.7 |
| 237CO | 237CO | 212.1 | 18.0 | 8.5 | 97.2 | 206.2 | 91.2 | 187.9 | 90.0 | 7.3 | 2.3 | 18.8 | 2.0 |
| 238CO | 238CO | 181.1 | 12.5 | 9.4 | 97.3 | 176.3 | 91.3 | 161 | 89.5 | 8.5 | 1.8 | 16.6 | 1.0 |
| 23CO | 23CO | 238.3 | 7.3 | 7.6 | 98.3 | 234.3 | 73.2 | 171.6 | 87.8 | 8.3 | 3.7 | 36.1 | 0.3 |
| 244C | 244C | 94 | 5.9 | 6.1 | 99.3 | 93.3 | 86.7 | 80.9 | 88.5 | 8.5 | 2.8 | 6.4 | 1.2 |
| 253C | 253C | 182.7 | 7.8 | 10.0 | 97.0 | 177.3 | 89.5 | 158.7 | 88.8 | 9.1 | 2.0 | 20.4 | 0.7 |
| 296C | 296C | 171.9 | 11.6 | 7.5 | 98.5 | 169.3 | 91.4 | 154.7 | 89.9 | 7.3 | 2.6 | 14 | 1.7 |
| 29I | 29I | 181.3 | 7.5 | 9.0 | 97.4 | 176.6 | 74.2 | 131.1 | 87.9 | 7.8 | 4.2 | 20.9 | 0.3 |
| 388C | 388C | 189.4 | 28.8 | 20.1 | 86.7 | 164.1 | 80.0 | 131.4 | 87.4 | 8.7 | 3.4 | 14.2 | 0.9 |
| 396C | 396C | 96.4 | 6.0 | 5.8 | 99.6 | 96 | 86.1 | 82.6 | 88.4 | 8.9 | 2.6 | 5.9 | 1.2 |
| 439C | 439C | 155.4 | 33.3 | 9.1 | 94.8 | 147.4 | 87.7 | 129.3 | 91.1 | 7.1 | 1.5 | 14.9 | 0.7 |
| 439C | 439C-b | 141.8 | 11.5 | 8.2 | 99.1 | 140.5 | 86.6 | 121.6 | 89.6 | 7.8 | 2.5 | 12.8 | 0.8 |
| 454C | 454C | 96.2 | 6.3 | 5.1 | 99.6 | 95.8 | 86.6 | 83 | 88.9 | 8.1 | 2.9 | 9.3 | 0.7 |
| 465C | 465C | 172.8 | 5.5 | 27.3 | 99.3 | 171.6 | 89.2 | 153.1 | 87.7 | 8.7 | 3.5 | 24.3 | 0.5 |
| 47I | 47I-a | 148.9 | 32.9 | 12.2 | 91.0 | 135.5 | 88.5 | 120 | 89.5 | 8.9 | 1.4 | 12.1 | 0.7 |
| 47I | 47I-b | 155 | 16.0 | 9.9 | 95.8 | 148.6 | 71.1 | 105.6 | 87.8 | 8.2 | 3.4 | 11.7 | 0.8 |
| 483C | 483C | 200.4 | 9.7 | 10.1 | 96.6 | 193.6 | 91.3 | 176.8 | 89.5 | 8.2 | 2.2 | 15.4 | 1.1 |
| 484C | 484C | 201.8 | 8.1 | 9.0 | 97.2 | 196 | 91.2 | 178.8 | 87.7 | 10.3 | 2.0 | 18.4 | 0.7 |
| 49I | 49I-a | 172.9 | 23.6 | 19.0 | 88.0 | 152.1 | 82.5 | 125.5 | 88.2 | 8.6 | 2.6 | 14.1 | 0.8 |
| 49I | 49I-b | 173.8 | 26.0 | 13.6 | 92.5 | 160.8 | 73.3 | 117.9 | 85.8 | 7.9 | 6.2 | 14.5 | 0.7 |
| 53I | 53I | 159.7 | 35.2 | 15.2 | 90.7 | 144.8 | 77.8 | 112.6 | 85.6 | 8.1 | 6.2 | 13.7 | 0.6 |
| 59I | 59I | 157.3 | 18.7 | 17.3 | 89.5 | 140.7 | 77.2 | 108.7 | 86.3 | 8.8 | 4.8 | 14.4 | 0.5 |
| 8CO | 8CO | 206.4 | 8.0 | 9.5 | 96.4 | 199 | 73.7 | 146.7 | 87.0 | 8.5 | 4.4 | 22.6 | 0.5 |
| 192C | 92C | 222.4 | 24.8 | 18.4 | 87.4 | 194.4 | 78.3 | 152.2 | 85.8 | 9.3 | 4.7 | 19.8 | 0.5 |

Technical summary of all libraries of this dataset. TSO: template switch oligo.

**Table S3.**

| Cells used for Plot | nGenes | nPCs | Neighbors | PAGA variable | PAGA res | Neighbors | Diffmap dimensions | Final UMAP spread |
|---|---|---|---|---|---|---|---|---|
| All Cells | 2500 | 56 | 100 | Cell type | 0.5 | 100 | 34 | 0.1 |
| Immune | 2000 | 28 | 50 | Cell type | 0.4 | 100 | 24 | 0.1 |
| Epithelial | 2000 | 20 | 50 | Cell type | 0.35 | 60 | 12 | 0.1 |
| Mesenchyme/Stromal | 2000 | 18 | 40 | Cell type | 0.2 | 50 | 13 | 0.1 |
| Classical Monocyte + Macrophage | 800 | 7 | 80 | Cell type | 0.2 | 120 | 5 | 0.5 |
| Fibroblast + Myofibroblast | 1500 | 25 | 10 | Louvain (res=0.7; seed=7) | 0.25 | 80 | 5 | 0.1 |

Values for all non-default parameters used during the generation of each UMAP.

**Table S4.**

| catalogue label | gene | clone | catalogue number | company | Host species |
|---|---|---|---|---|---|
| Cytokeratin 17 | KRT17 | E3 | V2176SAF-100UG | NSJ Bioreagents | mouse |
| CD31 | PECAM1 | JC/70A | MA513188 | Thermo Fisher Scientific | mouse |
| ITGA8 | ITGA8 | CL7304 | ab243027 | Abcam | mouse |
| COX2 | PTGS2 | COX 229 | 35-8200 | Thermo Fisher Scientific | mouse |
| Pro-Collagen I | COL1A1 | SP1.D8 | | Developmental Studies Hybridoma Bank | mouse |
| p16 | CDKN2A | EPR1473 | ab108349 | Abcam | rabbit |
| p21 | CDKN1A | EPR362 | ab109520 | Abcam | rabbit |
| p63 | TP63 | TP63/1423R | V3815-100UG | NSJ Bioreagents | rabbit |
| α-Smooth Muscle Actin | ACTA2 | D4K9N | 19245S | Cell Signaling Technology | rabbit |
| COL15A1 | COL15A1 | polyclonal | PA553667 | Thermo Fisher Scientific | rabbit |
| HMGA2 | HMGA2 | EPR18114 | ab207301 | Abcam | rabbit |
| CD34 | CD34 | EP373Y | ab81289 | Abcam | rabbit |
| Periostin | POSTN | EPR20806 | ab227049 | Abcam | rabbit |

Table of primary antibodies used in the immunohistochemical stainings.

**Data S1. (Data_S1_avgSubjectCelltype.markers.epiOnly.Final.txt)**
Results of Wilcoxon rank-sum test of each epithelial cell-type against the other epithelial varieties, using the average gene expression per subject, per cell type.

**Data S2. (Data_S2_avgSubjectCelltypeMarkers.mesenchymeOnly.Final.txt)**
Results of Wilcoxon rank-sum test of each mesenchymal cell-type against the other mesenchymal varieties, using the average gene expression per subject, per cell type.

**Data S3. (Data_S3_avgSubjectCelltypeMarkers.myeloidOnly.Final.txt)**
Results of Wilcoxon rank-sum test of each myeloid cell-type against the other myeloid varieties, using the average gene expression per subject, per cell type.

**Data S4. (Data_S4_avgSubjectCelltypeMarkers.lymphoidOnly.Final.txt)**
Results of Wilcoxon rank-sum test of each lymphoid cell-type against the other lymphoid varieties, using the average gene expression per subject, per cell type.

**Data S5. (Data_S5_AllCellTypesMarkers.logDOR.Final.txt)**
Log transformed diagnostics odds ratio of genes for cell types across cells, using genes with log transformed fold change > 0.3 for each cell population compared against the other cell populations

**Data S6. (Data_S6_CellCountsPerTypePerSubject_08152019.txt)**
Number of cells profiled from each described cell type, for each subject.

**Data S7. (Data_S7_IPFvCtrl.wilcoxDE.collapsed.08292019.txt)**
Results of Wilcoxon rank-sum test of average per subject gene expression, within each cell type comparing IPF to control. Cell types represented by less than 5 subjects in either disease category were not tested.

**Data S8. (Data_S8_COPDvCtrl.wilcoxDE.collapsed.08292019.txt)**
Results of Wilcoxon rank-sum test of average per subject gene expression, within each cell type comparing COPD to control. Cell types represented by less than 5 subjects in either disease category were not tested.

**Data S9. (Data_S9_IPFvCOPD.wilcoxDE.collapsed.08292019.txt)**

Results of Wilcoxon rank-sum test of average per subject gene expression, within each cell type comparing IPF to COPD. Cell types represented by less than 5 subjects in either disease category were not tested.
